# NOXA expression drives synthetic lethality to RUNX1 inhibition in pancreatic cancer

**DOI:** 10.1101/2021.10.21.465266

**Authors:** Josefina Doffo, Stefanos A. Bamopoulos, Hazal Köse, Felix Orben, Chuanbing Zang, Miriam Pons, Alexander T. den Dekker, Rutger W. W. Brouwer, Apoorva Baluapuri, Stefan Habringer, Maximillian Reichert, Anuradha lllendula, Oliver H. Krämer, Markus Schick, Elmar Wolf, Wilfred F. J. van IJcken, Irene Esposito, Ulrich Keller, Günter Schneider, Matthias Wirth

## Abstract

Evasion from drug-induced apoptosis is a crucial mechanism of cancer treatment resistance. The pro-apoptotic protein NOXA marks an aggressive pancreatic ductal adenocarcinoma (PDAC) subtype. To identify drugs that unleash the death-inducing potential of NOXA, we performed an unbiased drug screening experiment. In *NOXA*-deficient isogenic cellular models we identified an inhibitor of the transcription factor heterodimer CBFβ/RUNX1. By genetic gain and loss of function experiments we validated that the mode of action depends on RUNX1 and NOXA. Of note, RUNX1 expression is significantly higher in PDACs compared to normal pancreas. We show that pharmacological RUNX1 inhibition significantly blocks tumor growth *in vivo* and in primary patient-derived PDAC organoids. Through genome wide analysis, we detected that *RUNX1*-loss reshapes the epigenetic landscape, which gains H3K27ac enrichment at the *NOXA* promoter. Our study demonstrates a previously unknown mechanism of NOXA-dependent cell death, which can be triggered pharmaceutically. Therefore, our data show a novel way to target a therapy resistant PDAC, an unmet clinical need.

**Significance:** Recent evidence demonstrated the existence of molecular subtypes in pancreatic ductal adenocarcinoma (PDAC), which resist all current therapies. The paucity of therapeutic options, including a complete lack of targeted therapies, underscore the urgent and unmet medical need for the identification of targets and novel treatment strategies for PDAC. Our study unravels a function of the transcription factor RUNX1 in apoptosis regulation in PDAC. We show that pharmacological RUNX1 inhibition in PDAC is feasible and leads to NOXA-dependent apoptosis. The development of targeted therapies that influence the transcriptional landscape of PDAC might have great benefits for patients who are resistant to conventional therapies. RUNX1 Inhibition as a new therapeutic intervention offers an attractive strategy for future therapies.

## Introduction

Pancreatic ductal adenocarcinoma (PDAC) is an aggressive disease often diagnosed at an advanced stage. The incidence of PDAC is steadily increasing, and PDAC is predicted to be the second leading cause of cancer-related death by 2030 (1). Evasion of apoptosis is a characteristic of PDAC and is often associated with treatment resistance (2, 3). A dysregulated transcription commonly results in apoptosis resistance (4). Therefore, the identification of novel concepts to reactivate apoptosis by disrupting cancerous transcription programs is a promising approach for the effective elimination of PDAC cells (3, 5).

Comprehensive integrated genome analyses from RNA expression profiles in recent years revealed different subtypes of PDAC with variable biology and therapeutic responsiveness (6–9). *NOXA* (latin for damage; also known as PMAIP1 - Phorbol-12-myristate-13-acetate-induced protein 1) is part of an identifier gene-set for the quasi-mesenchymal subtype of the disease (8). This subtype overlaps with the described basal-like and squamous subtype of the disease, which is particularly resistant to the currently used chemotherapeutics (7, 9).

NOXA belongs to the BCL-2 homology (BH) BH3-only subgroup of the B-cell lymphoma 2 (BCL2) protein family which is essential for the regulation of cell intrinsic apoptosis (10). BCL2 proteins are divided into sensor, effector and protector proteins (10). The classical anti-apoptotic protector proteins including Myeloid cell leukemia 1 (MCL1) inhibit effector proteins (e.g. Bcl-2-associated X protein) thereby blocking apoptosis. Pro-apoptotic sensor proteins including NOXA, which directly binds to MCL1, neutralize the anti-apoptotic function of the protector proteins (10), leading to the initiation of apoptotic cell death. In PDAC, NOXA is tightly regulated at the transcriptional level, and transcriptional activation of *NOXA* by HDAC-inhibitors or proteasome inhibitors contribute to induce the cell intrinsic apoptosis pathway (11–13). Furthermore, *NOXA* is regulated by multiple transcription factors including p53 and is involved in apoptosis under genotoxic stress (14, 15).

Runt-related (RUNX) proteins are master regulators involved in a broad range of biological processes including proliferation, differentiation and apoptosis (16). DNA binding of these transcription factors is mediated by heterodimerization of a core DNA binding factor alpha chain (CBFα), composed of one of the three RUNX family members, RUNX1-3, to the non-DNA binding core binding factor beta (CBFβ). Each of the three RUNX family members play important roles in different stages of tumor development (17). As has been shown in mouse models, knockouts of any of the three RUNX transcription factors exhibit significant developmental defects: RUNX1 plays an important role in hematopoiesis (18), RUNX2 in bone development (19) and RUNX3 in the gastrointestinal tract (20) and in neurogenesis (21). RUNX expression patterns are highly dynamic and depend on the stage of differentiation, development and environmental conditions (22). In addition, RUNX transcription factors are expressed in almost all cancers (23). Besides its implication in leukemogenesis (24), RUNX1 is strongly expressed in a broad spectrum of solid tumors (25) and is associated with poor prognosis in PDAC (26). Depending on the cellular context, RUNX1 can act both oncogenic and tumor suppressive in solid tumors (27). RUNX1 interacts with various co-factors to shape gene expression. RUNX1-dependent activation of target genes is mediated through an interaction with CBP/p300 (28) and the protein arginine methyltransferase 1 (PRMT1) (29). The inhibitory function of RUNX1 is achieved by interaction with co-repressors such as the Sin3A-HDAC corepressor complex (30).

In this study, we aimed at identifying novel strategies affecting the delicate balance of NOXA expression to drive cell death in an aggressive subtype of PDAC. We found that inhibition of RUNX1 led to a global gain of H3K27ac enrichment contributing to the activation of the proximal *NOXA* promoter region and suggest a strategy to overcome treatment resistance in an aggressive subtype of PDAC with inferior prognosis.

## Materials and Methods

### Cell culture, cell viability assay

Cell lines were cultured in Dulbecco’s Modified Eagle’s Medium (Thermo Fisher Scientific, #41965062) or Roswell Park Memorial Institute (RPMI) 1640 Medium (Thermo Fisher Scientific, #21875091) supplemented with 10% FBS (Thermo Fisher Scientific, #10270106) and 1% Penicillin/Streptomycin (Thermo Fisher Scientific, #15070063). They were passaged up to 14 times in a 1:10 dilution every 3-4 days. Murine PDAC cell lines were generated from *Kras*^*G12D*^ driven mouse models as described (13). For all cell lines used, PCR-based mycoplasma tests were performed at regular intervals. Cell viability was measured by MTT-Test (Sigma-Aldrich, #M5655). Detailed information on the procedures cell viability, drug screening and colony formation assay in Supplementary Materials and Methods.

### Patient-derived organoids

PDAC biopsies and tissues were received from endoscopy punctures or surgical resection. 3D organoids were collected, propagated, and analyzed in agreement with the declaration of Helsinki. This study was approved by the ethical committee of TUM (Project 207/15). Written informed consent from the patients for research use of tumor material was obtained prior to the use. Detailed information in Supplementary Materials and Methods.

### *In vivo* drug efficacy analysis in mice, IHC

Xenograft assays were performed by EPO (Experimental Pharmacology and Oncology, Berlin-Buch). All animal experiments were approved by the local responsible authorities and performed in accordance with the German Animal Protection law. Detailed information in Supplementary Materials and Methods.

### Statistical Analysis, qChIP, ChIPseq, RNAseq, ATAC seq, 4C, qPCR, Western blot, CRISPR-editing

Detailed information on the procedures and data analyses in Supplementary Materials and Methods and in STable 5.

## Results

### High *NOXA* mRNA expression is associated with an aggressive PDAC subtype

As a stress response, tumor cells may express pro-apoptotic effectors that can be neutralized by anti-apoptotic counterparts, thus dampening the apoptotic response. Such tumor cells expose apoptotic potential. To investigate, if classical pro-apoptotic BH3-only proteins may contribute to this phenotype, we analyzed the mRNA expression of BH3-only genes (6) and filtered for transcriptome profiles of human PDAC tumors (Fig. 1A). From this data, we extracted indicated classical BH3-only genes, performed a hierarchical clustering based on the mRNA expression of the BH3-only genes and subdivided PDAC patient samples into the squamous subtype and combined the ADEX-, classical- and the immunogenic subtypes as “other” subtypes. We subsequently determined if the mRNAs of the listed classical BH3-only members are significantly enriched in these two groups. Of note, specifically high expression of *NOXA* was significantly (p < 0.001) associated with the squamous subtype (Fig. 1A and SFig1A), which is in line with the described expression in quasi-mesenchymal (QM) cancers (8). In accordance with this finding, high *NOXA* expression (>75^th^ percentile) characterized a PDAC patient cohort with inferior survival as compared to patients with low *NOXA* expression (<25^th^ percentile) in ICGC/Bailey et al. (Fig. 1B and SFig1B) and TCGA datasets (SFig. 1C). We hypothesized that the high expression of *NOXA* indicates that these tumors harbor an apoptotic potential, and that shifting the balance of apoptosis regulators to a pro-apoptotic state may constitute a therapeutic strategy.

**Figure 1.**
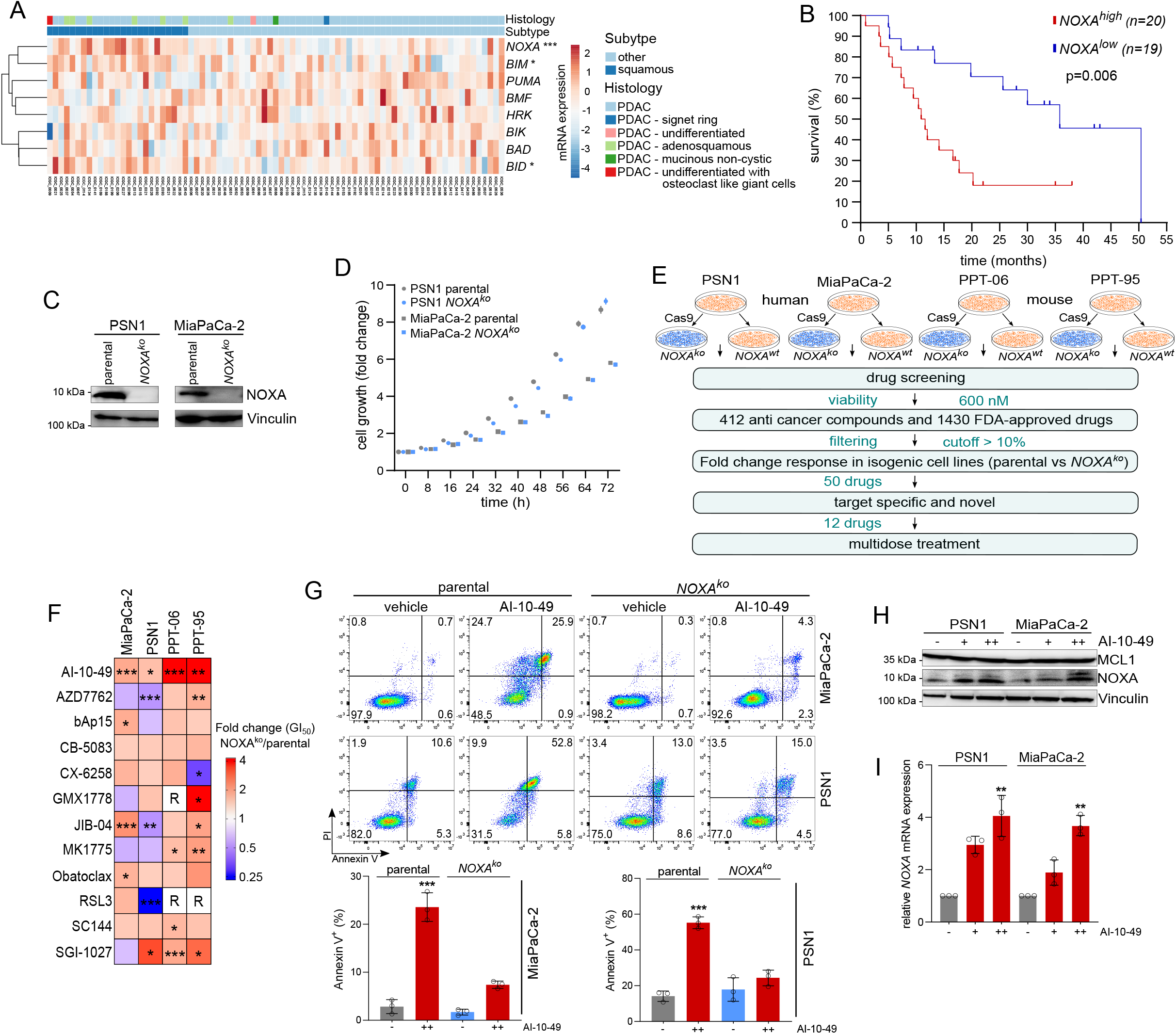
Screening for NOXA associated vulnerabilities in PDAC cells. **A)** Cluster analysis of mRNA from classical BH3-only proteins, derived from transcriptome profiles of PDAC patients (6). Molecular subtypes have been divided into two groups: the aggressive squamous subtype and all other subtypes have been merged to others. Histological Subtypes are indicated. Upper and lower quartiles of indicated mRNAs were identified. Significant (Fisher exact test) frequencies of a high mRNA expression of the indicated genes (upper quartile) in the squamous subtype are indicated (*p<0.05, **p<0.01, ***p<0.001). **B)** Survival of PDAC patients with a low (lower quartile) and a high (upper quartile) *NOXA* mRNA expression, derived from the dataset described in A)(6). Log rank test: p=0.006. **C)** Western blot analysis of NOXA protein in PSN1 and MiaPaCa-2 parental and isogenic *NOXA*^ko^ cell lines. Vinculin served as loading control. **D)** Growth curves of PSN1 and MiaPaCa-2 parental and isogenic *NOXA*^*ko*^ cell lines performed with live-cell imaging. 5 pictures per well were taken every 8 h and growth was calculated as confluence (%) normalized to 0 h control. **E)** Schematic representation of the performed high throughput drug screening strategy. 4 human pancreatic cancer cell lines (2 parental and 2 *NOXA*^*ko*^) and 4 murine cell lines (2 parental and 2 *NOXA*^*ko*^) were used for screening a total of 1842 drugs. These compounds were added to each cell line 24 h after seeding at a concentration of 600 nM and cell viability was measured by MTT assay after 72 h. n=3; all biological replicates were performed as technical triplicates. The inhibitors that differentially reduced viability in parental cell lines up to 10% more in comparison to *NOXA*^*ko*^ cells were further followed. Based on target treatment and or/novelty, 12 drugs were selected out of the first 50 hits. The GI_50_ of the drugs for murine and human cell lines was calculated from dose-response treatment using MTT assay. **F)** Dose-response treatment in 8 human and murine pancreatic cancer cell lines (4 parental, 4 *NOXA*^*ko*^). The fold change of the GI_50_ of the knockout cell lines compared to the parental is depicted. n=3; all biological replicates were performed as technical triplicates. Red represents sensitivity in parental cell line in respect to its isogenic counterpart (smaller GI_50_). Blue stands for higher sensitivity in the knockout cell line. R= resistant cell line within the used doses. Dose-response inhibition was calculated with logarithmic regression and tested for significance with logit model (*p<0.05, **p<0.01, ***p<0.001). **G)** FACS analysis of Annexin V/PI stained parental and *NOXA*^*ko*^ and cells after 72 h treatment with 3μM AI-10-49 (++) or DMSO (-) as vehicle control. n=3; all biological replicates were performed as technical triplicates. p value of unpaired t-test ***p<0.001. **H)** Western blot analysis of NOXA and MCL1 proteins in pancreatic cancer cell lines upon 6 h AI-10-49 treatment. Representative Western blot is shown. Vinculin served as loading control. n=3; all biological replicates were performed as technical triplicates. (-) DMSO, (+) 1.5 μM AI-10-49, (++) 3 μM AI-10-49. **I)** qPCR of *NOXA* in PSN1 and MiaPaCa-2 cell lines. Conditions as described in H).

### Identification of a synthetic lethal interaction of NOXA and inhibition of RUNX1

By analyzing transcriptome profiles, we selected human PDAC cell lines of the quasi-mesenchymal subtype (8) (SFig 1D) and Kras^G12D^-driven murine PDAC cell lines that exhibit relatively high basal *Noxa* mRNA expression (SFig. 1E) for cross-species validation to identify drugs that affect the apoptotic balance. To investigate vulnerabilities specifically created by NOXA expression, we generated human and murine isogenic PDAC cell line models with genetically defined NOXA status (SFig. 2A-D). To prove *NOXA* deficiency, we performed western blotting of the two human PDAC cell lines PSN1 and MiaPaCa-2. NOXA protein was absent in *NOXA*^*ko*^ cell lines (Fig. 1C). Importantly, *NOXA* deficiency did not influence the proliferation of the isogeneic cell lines (Fig. 1D).

To identify vulnerabilities associated with NOXA, we next performed a drug screening with a total of 1842 compounds in *NOXA-*proficient (parental) *and NOXA*-deficient (*NOXA*^*ko*^) cells and measured viability (Fig. 1E). Drug testing and viability assays were performed with a single concentration of 600 nM, as previously described (13). For identification of effects due of the NOXA status, we used a cut-off of 10% difference in viability (Fig. 1F). Out of the 1842 compounds we identified 50 drugs that showed higher efficiency in parental cell lines compared to *NOXA*^*ko*^ cell lines. Importantly, within the hits found, we identified topoisomerase and proteasome inhibitors, which is in line with data from previous studies (11–13), underlining the robustness of our screening experiment (STable 1). From our screening hits, we selected for specific targeted molecules (target specificity) and novelty (Fig. 1E) and further validated 12 hits with multi-dose experiments (Fig. 1F). Only one of these twelve compounds analyzed in the validation experiments, AI-10-49, showed no growth inhibition in *NOXA*^*ko*^ cell lines whereas viability of *NOXA*-proficient cells was dose-dependently reduced (Fig. 1F). AI-10-49 was originally designed to inhibit the interaction of the oncofusion protein CBFβ-SMMHC with RUNX1 (32). The lead compound of the bivalent AI-10-49 has been shown to inhibit RUNX1/CBFβ (33). Co-immunoprecipitations with either RUNX1 or CBFβ revealed AI-10-49 as RUNX1/CBFβ inhibitor in MiaPaCa-2 cells (SFig. 2E). Additionally, using the SwissTargetPrediction tool (34), CBFβ was a predicted target of this compound (STable 2). In summary, our data suggest that impairment of RUNX1 activity may affect NOXA-dependent execution of cell death in PDAC.

### Induction of NOXA by CBFβ/RUNX1 inhibition

To investigate whether the AI-10-49-induced drop in viability is mediated by induction of the apoptotic process, we performed fluorometric analysis of Annexin V/PI stained PDAC cells. Indeed, the reduced viability in parental cell lines upon AI-10-49 treatment was clearly associated with a significant induction of cell death in parental cells, whereas only marginal apoptosis induction was observed in *NOXA*^*ko*^ cells (Fig. 1G).

We next investigated whether an altered expression of other BCL2 family members such as MCL1, BCL2, BCLx_L_, BIM, BID, BAK or BAX mediates AI-10-49-induced apoptosis. We detected PARP cleavage but did not detect differential expression of these BCL2 family proteins 6 hours and 24 hours after AI-10-49 treatment (Fig. 1H, SFig. 3A, B), a marked induction of NOXA protein (Fig. 1I) and *NOXA* mRNA expression (Fig. 1I) was observed upon treatment with AI-10-49. In HCT116 cells harboring wild-type p53, a DNA damage stimulus induced both RUNX1 and p53 and activated p53 target genes, including *NOXA* (35). To test if p53 is involved in AI-10-49 induced *NOXA* expression we treated murine cell lines harboring either wiltype, mutant or deleted p53- (SFig. 3C) with AI-10-49 and the topoisomerase II inhibitor etoposide (SFig. 3D). While *Noxa* induction was highest in wildtype p53 cells upon etoposide treatment, we observed a rather uniform induction of *Noxa* in each of these cell lines after AI-10-49 treatment (SFig. 3D). Since p53 is mutated in the human PDAC cell lines PSN1 and MiaPaCa-2 we also analyzed the expression of the p53 family member p63, but did not detect significant regulation (SFig. 3E). Both p63 and mutant p53 showed no difference in expression at early time points after treatment with AI-10-49, but tended to show decreased expression after 48 h and 72 h, respectively (SFig. 3B,E). Our data show that both p53^mut^ and RUNX1 are not induced upon AI-10-49 treatment (SFig. 3B, E). Contrary, we rather observed a decreased expression of RUNX1 after AI-10-49 treatment in MiaPaCa-2 and PSN1 cells (SFig. 3E).

Next, to substantiate caspase-induced apoptosis, we examined AI-10-49 together with the pan-caspase inhibitor zVAD-FMK by Annexin V/PI FACS. Here, we observed a significant rescue in apoptosis induction (Annexin V^+^/PI^−^ fraction) (SFigure 3F). Both fractions, i.e. Annexin V^+^/PI^−^ and Annexin V^+^/PI^+^, remained unaffected in *NOXA*^*ko*^ cells, indicating the relevance of NOXA in cell death (SFigure 3F).

To further investigate the role of NOXA in AI-10-49 induced cell death, we generated MiaPaCa-2 cells stably expressing the CRISPR activator (CRISPRa) dCas9-MS2-p65-HSF1 (MpH), and an established *NOXA* sgRNA to endogenously overexpress NOXA (SFig. 2E, SFig. 3G). *NOXA*-CRISPRa cells phenocopied AI-10-49 treated cells in clonogenic assays and, more importantly, *NOXA*-CRISPRa cell growth was drastically inhibited when treated with AI-10-49 (SFig. 3H, I).

Together, our data show that inhibition of RUNX1 by AI-10-49 induces *NOXA* mRNA and protein expression and thereby drives NOXA-dependent apoptosis.

### RUNX1 is upregulated in pancreatic cancer and suppresses *NOXA* expression

Understanding the mode of action of drugs is critical for the implementation of patient stratification strategies. To address this question, we next investigated how AI-10-49 induces *NOXA* expression. Since AI-10-49 inhibits the interaction between CBFβ and a DNA binding α subunit encoded by RUNX proteins (16), we tested whether loss of RUNX expression affects NOXA expression in knockouts of all three *RUNX* genes *RUNX1*, *RUNX2* and *RUNX3* in MiaPaCa-2 cells (SFig 2F). Remarkably, we observed an induction of *NOXA* mRNA solely in *RUNX1* knockout cells (Fig. 2A), arguing for a RUNX1-specific repression of the *NOXA* gene. We validated this effect by siRNA-mediated *RUNX1* depletion in Panc1, AsPC1 and MiaPaCa-2 cells (SFig. 4A). In addition to the induction of *NOXA* mRNA, we also observed a significant NOXA induction in *RUNX1*^*ko*^ cells at the protein level (Fig. 2B). To analyze whether *RUNX1*^*ko*^ cells have a growth disadvantage, we compared colony formation of *RUNX1*^*ko*^ cells to the parental cell line. Here, we observed significantly reduced formation of colonies (Fig. 2C), which phenocopied the effects observed in AI-10-49 treated cells (SFig. 3H, I). In addition, *RUNX1*^*ko*^ cells showed an increased cell death rate at basal levels, as demonstrated by Annexin V flow cytometry (Fig 2D). To demonstrate the specificity of AI-10-49 in this context, we applied AI-10-49 in two *RUNX1*^*ko*^ clones as well as in *RUNX1* siRNA-treated MiaPaCa-2 cells. Although we did not observe a consistent effect in the *RUNX1* knockout clones, which is probably related to the difficult cultivation of these cells, the use of siRNA, on the other hand, showed reproducible effects equivalent to a doubling of the GI_50_ value, confirming the dependence on RUNX1 (SFig. 4B). These genetic experiments identify RUNX1 as negative regulator of *NOXA* gene expression.

**Figure 2.**
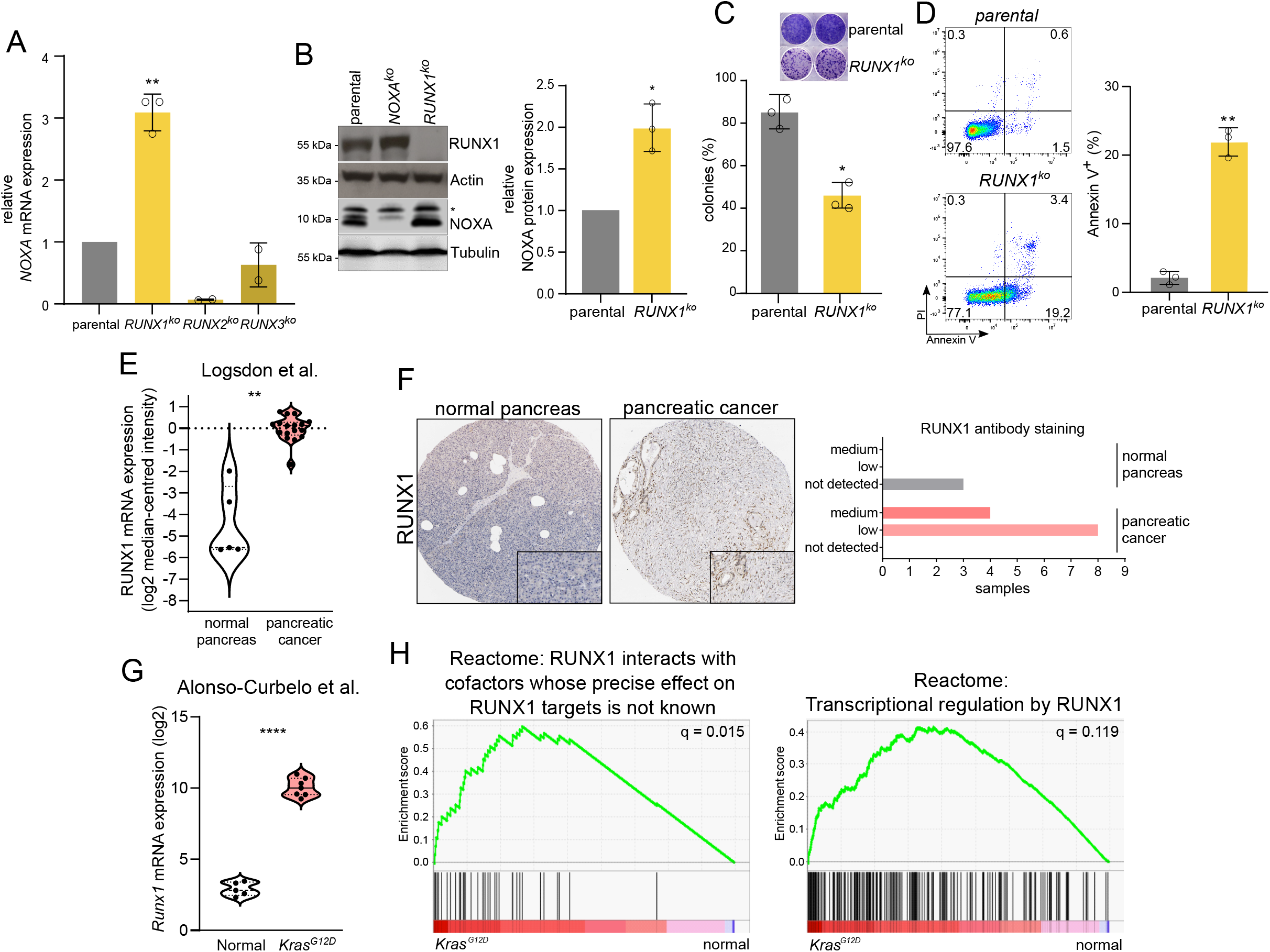
RUNX1 is upregulated in pancreatic cancer and a genetic RUNX1 deletion induces NOXA transcription and apoptosis. **A)** qPCR of *NOXA* in parental, *RUNX1*^*ko*^, *RUNX2*^*ko*^ and *RUNX3*^*ko*^ MiaPaCa-2 cells. mRNA fold change was calculated in comparison to parental cell line. n=3; all biological replicates were performed as technical triplicates. **B)** Western blot analysis and quantification of NOXA and RUNX1 proteins in parental, *NOXA*^*ko*^ and *RUNX1*^*ko*^ MiaPaCa-2 cells. Tubulin and actin served as loading controls. * unspecific band. P value of t-test, p<0.05. **C)** Representative image and quantification of clonogenic assay in parental and *RUNX1*^*ko*^ cells. n=3; all biological replicates were performed as technical triplicates. P value of t-test, p<0.05. **D)** FACS analysis of Annexin V stained parental and *RUNX1*^*ko*^ MiaPaCa-2 cells. n=3; all biological replicates were performed as technical triplicates. P value of t-test, p<0.01. **E)** *RUNX1* mRNA expression of patient samples from normal pancreas and pancreatic cancer (50), accessed via the oncomine.org, p<0.01. **F)** IHC of patient samples from normal pancreas and pancreatic cancer accessed via the human protein atlas (51). **G)** *RUNX1* mRNA expression of mouse samples from normal pancreatic epithelial cells and Kras^G12D^ pancreatic epithelial cells, p<0.0001. **H)** Gene set enrichment analysis (GSEA) of dataset, described in (52) F).

To further investigate the relevance of RUNX1 in PDAC, we analyzed RUNX1 expression in human and murine datasets. We found that RUNX1 indeed is transcriptionally upregulated in pancreatic cancer in all three datasets investigated (Fig. 2E, SFig4C). In addition, RUNX1 was expressed in pancreatic cancer, whereas it was not detected in normal pancreas on protein level (Fig. 2F). In premalignant Kras^G12D^ driven murine pancreatic epithelial cells *Runx1* mRNA (Fig. 2G) and RUNX1 target gene signatures (Fig. 2H) were significantly enriched compared to normal pancreatic epithelial cells, displaying specificity for RUNX1 expression in non-stromal tumor-initiating cells. During PDAC progression RUNX1 expression is maintained. Suspecting a RUNX1-mediated repression of the *NOXA* gene, we re-analyzed the ICGC/Bailey et al. (6) and Collisson et al. (8) transcriptome datasets and indeed observed a negative correlation trend of *RUNX1* and *NOXA* expression in the squamous/quasi-mesenchymal PDAC subtype (SFig. 4D). To investigate whether RUNX1 expression was the main contributing factor to NOXA expression, we performed a shRNA-mediated knockdown of *NOXA* in *RUNX1*^*ko*^ cells (SFig. 4E). Congruent with our initial findings, the basal apoptosis rates were restored (SFig. 4F).

Taken together, these data argue for a RUNX1-mediated repression of NOXA expression.

### RUNX1 inhibition induces *NOXA* through amplification of active chromatin marks

To understand the immediate effect of AI-10-49 treatment, we performed RNAseq analyses of MiaPaCa2 cells 6 h upon treatment in comparison to vehicle treated cells. Apart from NOXA, which was upregulated in treated cells, less than 300 genes, were found to be differentially expressed, suggesting a high specificity of AI-10-49 (Fig. 3A). Gene set enrichment analyses revealed a significant apoptosis signature and a negative enrichment score for RUNX1 targets (Fig. 3B, STable 3) upon AI-10-49 treatment. To analyze the impact of RUNX1 inhibition on chromatin accessibility we performed assays for transposase accessible chromatin with high-throughput sequencing (ATAC-seq). We observed reduced chromatin accessibility after AI-10-49 treatment, compared to the vehicle control 6 h after AI-10-49 treatment (Fig. 3C). In addition, a comparison of parental MiaPaCa-2 cells with isogenic *RUNX1* deficient cells showed a an increased accessibility of the chromatin.

**Figure 3.**
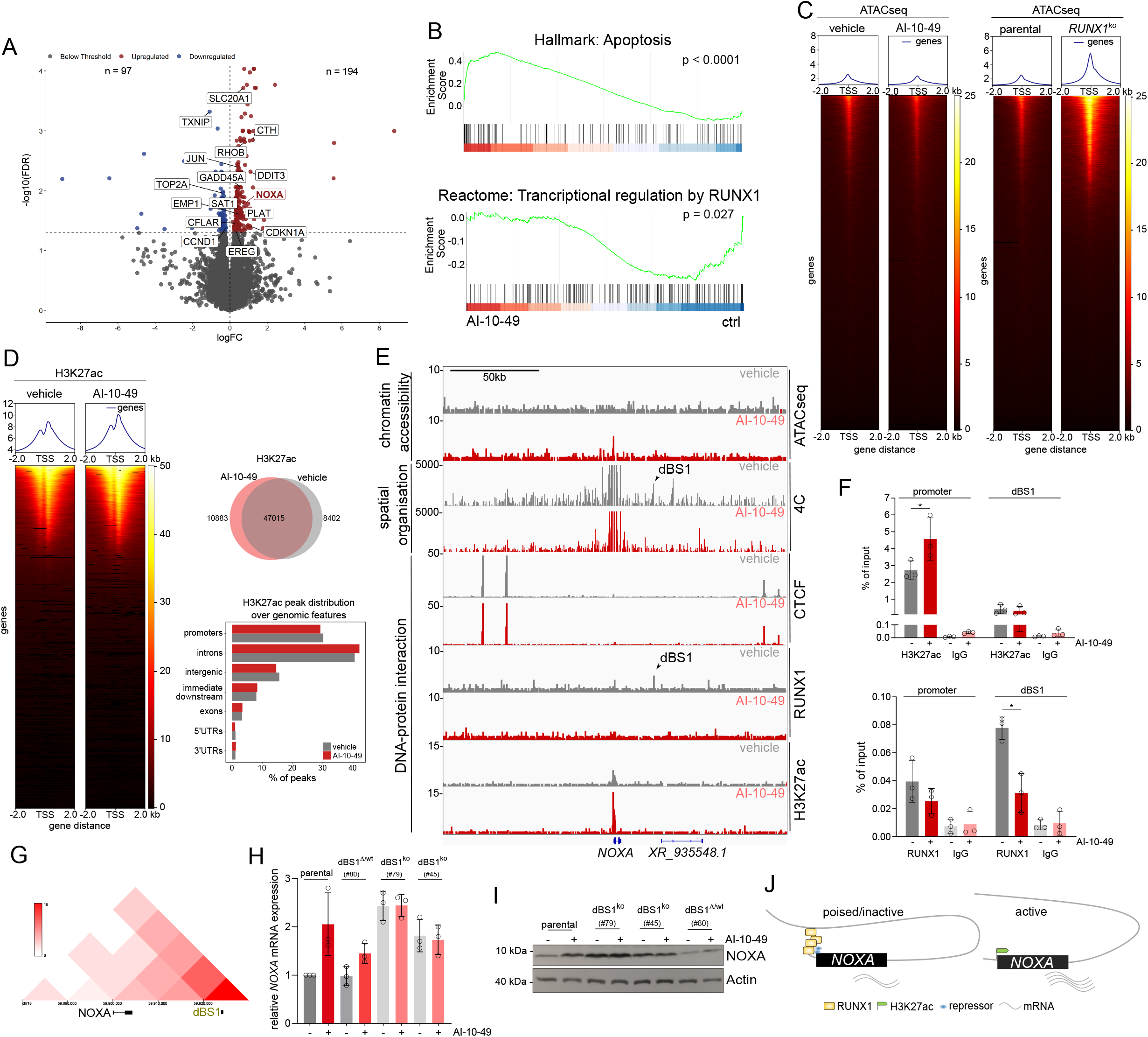
RUNX1 inhibition induces a global redistribution of active chromatin and activates of the proximal promotor region of the NOXA gene. **A)** Volcano Plot of differential expressed genes. MiaPaCa-2 cells were treated with vehicle (DMSO) or AI-10-49 (3μM) for 6 h (n=3 biological replicates). **B)** Indicated gene signatures of a GSEA from RNAseq data, described in A). GSEA, gene set enrichment analysis. **C)** Heatmap representing open chromatin peaks by omni-ATAC-seq analysis in DMSO and AI-10-49 treated cells (3μM for 6h) and in parental and *RUNX1*^*ko*^ MiaPaCa-2 cells (n=2). **D)** Heatmap representing H3K27ac peaks in MiaPaCa-2 cells treated for 6 h with vehicle (DMSO) or 3 μM AI-10-49 (n=2). Venn diagram displays H3K27ac peaks in AI-10-49 and vehicle treated MiaPaCa-2 cells. H3K27ac peak distribution over genomic features are displayed in a bar plot diagram. **E)** Omni-ATACseq (DNA accessibility) (n=2), chromosome conformation capture (4C, spatial chromatin organisation) (n=1) and ChIPseq (n=2) analysis for RUNX1, H3K27ac and CTCF in vehicle (DMSO)- or AI-10-49 treated MiaPaCa-2 cells. Arrows indicate either RUNX1 binding and an interaction of the *NOXA* gene at downstream binding site 1 (dBS1). **F)** ChIP-qPCR analysis at the promoter and the downstream binding site 1 (dBS1) for RUNX1, H3K27ac and IgG in DMSO- or AI-10-49 treated MiaPaCa-2 cells (n=3). **G)** Heatmap of Hi-C data from Panc1 cells generated by Dekker laboratory (36), accessed via http://3dgenome.fsm.northwestern.edu/ displaying spatial chromatin organization. **H)** Relative *NOXA* mRNA expression in indicated clones of MiaPaCa-2 cells. The dBS1 binding site was excised using CRISPR/Cas9. Cells were treated for 6h with 3μM of AI-10-49. **I)** Western blot of NOXA 6h upon treatment with 3μM AI-10-49 in indicated clones of MiaPaCa-2 cells. Actin served as loading control. **J)** Schematic of RUNX1 mediated repression of the *NOXA* gene.

To investigate the association of RUNX1 inhibition with regulation of chromatin dynamics, we performed H3K27ac ChIPseq experiments to detect transcriptionally active chromatin. A cross-coverage and fingerprint plot showed adequate signal strength in enriched regions (SFig. 5A, B). Globally, H3K27ac signal was increased in both replicates (Fig. 3D, SFig. 5C), arguing for a neutralization of the repressor activity of RUNX1.

We also performed RUNX1 ChIPseq to assess the impact of RUNX1 inhibition on RUNX1 binding. This analysis showed a peak downstream of the *NOXA* gene, which we hypothesized to be an enhancer region and coined it downstream binding site 1 (dBS1). To substantiate our findings, we analyzed RUNX1 ChIPseq from K562 and MCF10A cells. Indeed, we found an overlap of our identified peak in both ChIPseq datasets and were thus able to validate the peak identified in MiaPaCa-2 cells (SFig. 6A). In both, ChIPseq and qChIP experiments, we observed a drop in RUNX1 binding at downstream binding site 1 (dBS1) and an increased acetylation of H3K27 at the *NOXA* promoter upon AI-10-49 treatment (Fig. 3E, F), indicating an activation of the gene by acetylation of the proximal promoter region of *NOXA*, subsequently leading to increased gene expression. To analyze the spatial organization of the *NOXA* region, we performed chromosome conformation capture assays (4C) to capture interactions between the *NOXA* locus (view point) and all other genomic loci. Here, we found a hitherto unknown interaction with a downstream region of the *NOXA* gene, which is abrogated upon AI-10-49 treatment and in *RUNX1*^*ko*^ cells (Fig. 3E, arrow/dBS1, SFig. 6A). Binding of the nuclear protein CCCTC-binding factor (CTCF), which marks insulator regions to prevent crosstalk between active and inactive chromatin, was unaffected (Fig. 3E). Additionally, RUNX1 peaks in the vehicle control (indicated by an arrow in Fig. 3E) and at the *NOXA* gene, arguing for a spatial interaction, which is mediated by RUNX1 (Fig. 3E, G). The dBS1 region was the only region within the CTCF boundaries where both RUNX1 binding and DNA-DNA interaction had disappeared after AI-10-49 (Fig. 3G, SFig. 6A). This spatial interaction could also be found in public Hi-C data of Panc1, K562 cells, and to a lower extend in the epithelial cell line MCF10A (Fig. 3F, SFig. 6B) (36). Taken together, these data suggest that RUNX1 binding to the dBS1 region actively represses the *NOXA* gene. To identify which histone deacetylases are responsible for this effect, we used class I HDAC inhibitors. Here, in particular, inhibition of HDAC1/2 by Merck60 showed a significant induction of *NOXA* mRNA expression (SFig. 6C) as well as an induction of the H3K27ac mark at the *NOXA* promoter (SFig. 6D). Additionally, murine PDAC cells harboring a dual recombinase system and a 4-hydroxy-tamoxifen inducible Cre to knockout alleles for either *Hdac1* (PPT-F3641) or *Hdac2* (PPT-F1648) (SFig. 6E), displayed a *Noxa* induction only in *Hdac2* deleted cells (SFig. 6F), which is in line with a previous study showing that HDAC2 is responsible for the repression of *NOXA* in PDAC (11). Therefore, a HDAC2-RUNX1/CBFβ axis might be responsible for *NOXA* repression. This requires further validation.

To prove that the *dBS1* region is causative for the repression of *NOXA*, we first screened this region for evolutionary conserved *RUNX1* binding motifs, and indeed identified conserved RUNX1 consensus sequences (SFig.6G). We next performed a CRISPR/Cas9-mediated knockout of the *dBS1* region (SFig. 6H) to genetically demonstarte its involvement in NOXA repression. Indeed, loss of the *dBS1* region led to increased *NOXA* mRNA (Fig. 3H) and protein levels (Fig. 3I). This demonstrates the repressive function of this RUNX1 binding site. In contrast to parental cells, AI-10-49 treatment did not further affected NOXA expression (Fig. 3H, I). Furthermore, in the *dBS1*^*Δ/wt*^ clone NOXA expression still was induced upon AI-10-49 treatment, albeit to a lesser extent (Fig. 3H, I).

In summary, we describe a previously unknown mechanism of a RUNX1-mediated repression of the *NOXA* gene in PDAC. Through enrichment of an active chromatin mark at the *NOXA* gene itself, its expression is significantly increased (Fig. 3J), thereby inducing apoptosis. This mechanism could be crucial for therapeutic interventions that depend on a NOXA-induced cell death program.

### RUNX1 inhibition by AI-10-49 is effective *in vivo* and in patient derived organoids

To validate whether RUNX1 inhibition could be effective *in vivo*, we first examined the efficacy of AI-10-49 in mice carrying MiaPaCa-2 PDAC xenografts (Fig. 4A). AI-10-49 treatment resulted in a significant decrease in tumor volume (Fig. 4B) and proliferative capacity (Ki67, Fig. 4C). Importantly, AI-10-49 treatment induced apoptosis measured by cleaved caspase 3 (CC3) positivity in the tumor compared to the vehicle control in parental cells (Fig. 4B, C). Whereas parental cells did not change in tumor volume after AI-10-49 treatment, *NOXA*^*ko*^ cells still grew upon treatment, supporting NOXA as an essential contributor of AI-10-49 efficacy (Fig. 4B). Additionally, no significant difference between control and treatment was observed in either K67 or CC3 stainings in *NOXA*^*ko*^ cells (Fig. 4D). Taken together, these data indicate that apoptosis induction by AI-10-49 treatment is also dependent on NOXA *in vivo*.

**Figure 4.**
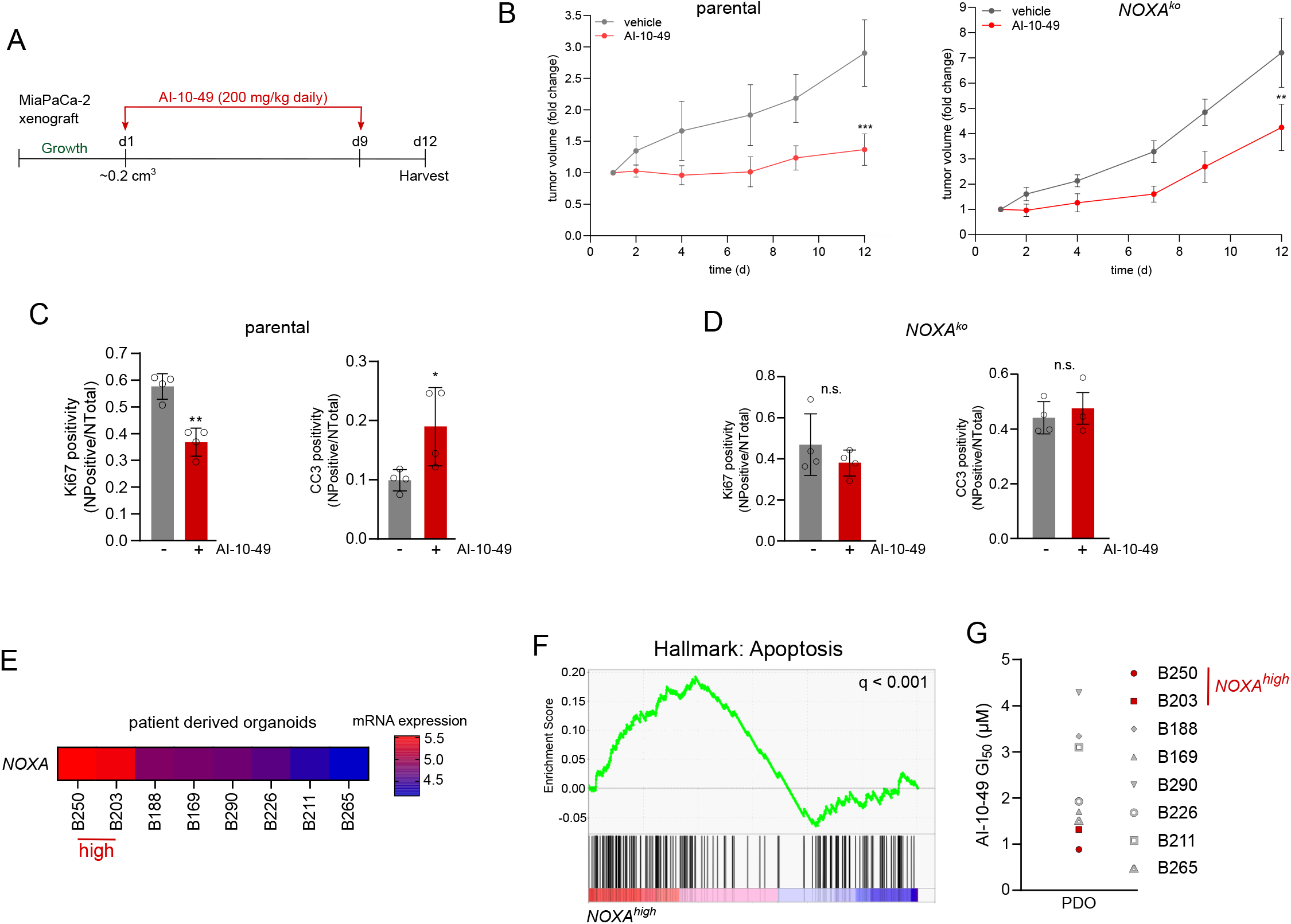
Tumor growth is blocked by RUNX1 inhibition in vivo and in patient derived organoids. **A)** Mice were treated with 200 mg/kg AI-10-49 intra peritoneal daily for 9 days. Treatment started (d1) when tumors reached a volume of 0.2 cm^3^. **B)** Tumor size was measured over time in parental and *NOXA*^*ko*^ xenografts. AI-10-49 treated mice showed a significant tumor growth inhibition (n=5 mice in each group).**p<0.01, ***p<0.001 (Student’s t-Test) **C)** IHC of parental xenografts. Left panel: Representative pictures of tumors from AI-10-49 and vehicle treated mice. Displayed are full scans of the tumors (bar: 2mm); detailed pictures of H&E stained and IHC for Ki67 and cleaved caspase 3 (CC3, bar: 100μm). Right panel: Quantification of Ki67 and CC3 IHC staining of AI-10-49 treated tumor xenografts using the Aperio positive pixel method. * p < 0.055, ** p < 0.01 (t-test). **D)** IHC of *NOXA*^*ko*^ xenografts as indicated in C). **E)** RNA-seq data of 7 patient derived organoids (PDOs) were analyzed for *NOXA* expression. *NOXA* mRNA expression > 75% = *NOXA*^*high*^; NOXA mRNA expression < 25% = *NOXA*^*low*^. **F)** GSEA of RNA-seq data of PDOs. Hallmark apoptosis signature in the *NOXA*^*high*^ subtype. Nominal p-value < 0.001. FDR-q value is depicted. GSEA, gene set enrichment analysis. **G)** Dose-response treatment of PDOs viability measured after 72h upon AI-10-49 treatment with CellTiterGlo^®^.

We next isolated seven human patient-derived organoids (PDOs) from PDAC patients to investigate RUNX1 inhibition by AI-10-49. First, we performed transcriptome profiling (STable 4), and sorted PDOs for high and low *NOXA* mRNA expression (Fig. 4E). Gene set enrichment analysis revealed a significant (q<0.001) accumulation of an apoptosis signature in the PDOs with a high *NOXA* expression (Fig. 4F). PDOs with a high *NOXA* expression showed the strongest growth inhibition towards AI-10-49 treatment, which further supports our previous findings (Fig. 4G).

Overall, these findings show that RUNX1 inhibition might be a novel therapeutic option to treat PDAC.

## Discussion

Molecular tumor profiling and functional studies have led to the identification and validation of genes and signaling pathways that are dysregulated or mutated in PDAC (7, 37–39). Based on comprehensive molecular characterization of PDACs (6–9), it might thus be possible to define personalized treatment strategies (38). An imbalance of signaling pathways, such as cell death-associated pathways, promote tumor maintenance and treatment resistance in PDAC (3). In this study, we analyzed publicly available transcriptome profiles of PDAC patients and found that increased *NOXA* mRNA expression defines an aggressive squamous/quasi-mesenchymal subtype. In contrast to NOXA, which is tightly regulated at the transcriptional level in PDAC (11), one of its anti-apoptotic counterparts, MCL1 (10), is mostly regulated at the protein level (40). Since *NOXA* mRNA and protein expression do not correlate strongly, as has been shown in mantle cell lymphoma (41) and in PDAC (11), it is important to identify substances that can induce apoptosis by taking advantage of a NOXA-associated vulnerability. We therefore performed drug-screening experiments in isogenic cell models with *NOXA*-deficient and *NOXA*-proficient counterparts to search for compounds that can exploit this vulnerability. We unexpectedly found a substance that inhibits the core binding alpha units RUNX1, RUNX2 or RUNX3 with CBFβ.

The functions of RUNX1 are highly specific depending on the tissue and cell type. Deletion of *Runx1* in a mouse model of T-cell acute lymphoblastic leukemia (T-ALL)(42), silencing of *RUNX1* in human T-ALL cells (42) and silencing of *RUNX1* in SW480 human colon cancer cells (43) all triggered apoptosis. In contrast, in Kasumi-1 t(8;21) leukemia cells RUNX1 overexpression induced apoptosis by eliciting expression of the cyclin-dependent kinase inhibitor p57^Kip2^ (44). In line with the biological effects observed in T-ALL (42) and in colon cancer cells (43), we observed an induction of apoptosis both through the pharmacological CBFβ/RUNX1 inhibition by the compound AI-10-49 and in CRISPR/Cas9 mediated knockouts of *RUNX1*. We observed a significant induction of *NOXA* mRNA expression, which was exclusive to *RUNX1* knockout cells and could not be observed in *RUNX2* or *RUNX3* knockout cells, arguing for a non-redundant function for RUNX1.

In a chemical high-throughput screen, which was performed to identify compounds that disrupt the interaction between RUNX1 and the CBFβ-MYH11/SMMHC fusion protein first a lead molecule was identified, which exhibited low selectivity for the fusion protein (33). Therefore, a bivalent derivate of this compound was generated. AI-10-49 inhibits CBFβ-MYH11/SMMHC with an increased selectivity and restores the formation of wild-type CBFβ-RUNX1 (33). In cells, lacking the CBFβ-MYH11/SMMHC fusion protein, AI-10-49 acts like the monomeric lead molecule and inhibits wild-type CBFβ/RUNX1. In fact, both AI-10-49 treatment as a putative pharmacological CBFβ/RUNX1 inhibitor, as well as a genetic knockout of *RUNX1* unexpectedly showed an induction of NOXA, similarities in transcriptional regulation at a genome-wide scale and an associated apoptosis induction in PDAC cells, arguing for RUNX1 as a repressor of *NOXA* gene expression. We could show that the pharmacological CBFβ/RUNX1 inhibition leads to a global enrichment of H3K27ac, a marker for active chromatin, including the *NOXA* gene. We observed an unexpected punctual interaction of a NOXA downstream RUNX1 binding site and the NOXA promoter. How exactly RUNX1 exercises its repressive function in PDAC has to be addressed in detail in further studies. An analysis of different data sets showed a high expression of RUNX1 in pancreatic cancer compared to normal pancreas. Of note, knockdown of *RUNX1* in PDAC cells was shown to suppress the invasive/aggressive phenotype via regulation of miR-93 (45). In particular, high RUNX1 expression and RUNX1 target gene signatures were observed in premalignant Kras^G12D^ driven murine pancreatic epithelial cells, which together indicate a largely unexplored and possibly unexploited relevance of RUNX1 in PDAC.

Since the apoptosis machinery in PDAC cells retains its functionality (3), the strategy of directly inhibiting pro-survival BCL2 proteins such as the NOXA antagonist MCL1 appears extremely attractive (46). One of the first selective MCL1 inhibitors with *in vivo* activity, S63845, showed massive apoptosis induction in multiple myeloma and acute myeloid leukemia, but many solid tumors were resistant to S63845 monotherapy (47). Combination therapies such as S63845 with the SRC kinase inhibitor Dasatinib reduced cell viability in PDAC models and even lead to a reduction in metastasis formation (48). This data shows that the development of new therapeutic strategies, especially with regard to apoptosis evasion for PDAC, are extremely promising in improving the current clinical regimens. Whether compounds like S63845 synergistically combine with AI-10-49 is currently under investigation.

Unraveling the transcriptional regulation of apoptosis-associated genes and the interplay of transcription factors, such as RUNX1, is important to understand the tumor biology of PDAC. The development and improvement of compounds that can inhibit transcription factors, such as the CBFβ/RUNX1 inhibitor used here, perhaps in combination with proteolysis-targeting chimeras (PROTAC) technology (49), could provide new approaches for PDAC treatment. Therefore, our mechanistic work, demonstrating a control of NOXA by a repressive facet of RUNX1 opens a novel research direction into potent RUNX1 inhibitors and a novel way to target this deadly disease.

## Acknowledgements

We thank Martin Schlensog and Yakup Yasar for performing the IHC staining. We thank Dieter Saur for providing murine PDAC cell lines and thank Ernesto Acevedo for providing the p53(1C12) antibody. Funding: Else-Kröner-Fresenius Stiftung (2016_A43 to M.W.); Walter Schulz Stiftung to M.W.; Deutsche Forschungsgemeinschaft (DFG) [SFB1321 (Project-ID 329628492) C13 to G.S. and SCHN 959/6-1 and SCHN 959/3-2 to G.S.] and SFB 1335 project P3 to U.K.; Wilhelm-Sander-Stiftung (2017.048.2 to G.S. and U.K.); Deutsche Krebshilfe project 70114425 to U.K.; Stiftung Charité to U.K..

## Availability of Data and Materials

RNAseq (cell lines), ChIPseq, ATACseq and 4C accession No.: PRJEB39828. RNAseq (organoids) gene expression matrix is shown in the STable 4.

## Supplements

**SFigure 1** Classical BH3 only mRNA expression and survival analysis in PDAC

**A)** mRNA expression of BH3-only differentially expressed in PDAC subtypes classified in squamous and other. Fisher’s exact test, * p < 0.01, ** p < 0.001, *** p < 0.0001. **B)** Simple Cox Regression models were constructed using the expression of all “classical BH3 family members”. Genes with p-values < 0.1 were used to construct a multiple Cox regression model. Out of all members, only *NOXA* significantly correlated with inferior survival (HR = 1.9, p < 0.001), while increasing expression of *BBC3/PUMA* was associated with higher overall survival ((HR = 0.73, p = 0.095). **C)** Survival of PDAC patients with a low (lower quartile) and a high (upper quartile) *NOXA* mRNA expression derived from curated TCGA datasets of PDAC patients as described (13). Log rank test as indicated. **D)** Human PDAC cell line subtypes according to Collisson et al. (8). QM: quasi-mesenchymal.

**SFigure 2** *CRISPR/Cas9 knockout and CRISPRa strategies and co-immunopreciptitation of RUNX1 and CBFb in AI-10-49 treated MiaPaCa-2 cells*

**A)** Scheme of the human *NOXA* gene (parental and knockout) with indicated primer binding sites. **B)** Genotyping PCR of indicated human PDAC cells to screen for a *NOXA* knockout. **C)** Scheme of the murine *NOXA* gene (parental and knockout) with indicated primer binding sites. **D)** Genotyping PCR of indicated murine PDAC cells to screen for a *NOXA* knockout. **E)** Co-immunoprecipitation of RUNX1 and CBFβ in MiaPaCa-2 cells. Cells were treated for 6h with 3μM of AI-10-49 or DMSO as vehicle control. Either a RUNX1 or a CBFβ antibody was used for the Co-IP as indicated. An IgG pulldown was used as negative control. **F)** Schematic representation of dCas9-VP64-MS2-HSF1 driven NOXA overexpression. One sgRNA drives dCas9 towards the *NOXA* promoter region to induce its expression. **G)** Schematic representation of RUNX gene family knockout. For RUNX1 knockout, 2 sgRNAs were designed to excise the gene region from exon 3 to exon 4. For RUNX2 knockout, 2 sgRNAs were designed to target excision of beginning of exon 8 until stop codon. For RUNX3 knockout, 2 sgRNAs target upstream exon 1 and end of exon 2.

**SFigure 3** *RUNX1 inhibition by AI-10-49 induces NOXA associated apoptosis*

**A)** Western blot analysis of indicated BCL2 family members and cleaved PARP upon 6h and 24h treatment with 3μM of AI-10-49 in MiaPaCa-2 and PSN1 cells. Actin served as loading control. **B)** Protein expression analysis of MCL1, p53, BCL-xL, BIM and BAX upon 24h and 72h treatment with 3μM of AI-10-49 and the pan-caspase inhibitor zVAD-FMK in parental and *NOXA*^*ko*^ MiaPaCa-2 cells. GAPDH served as loading control. **C)** p53 Western Blot of indicated murine PDAC cell lines to determine basal p53 expression of n=3 biological replicates. Actin served as loading control. **D)** Relative *Noxa* mRNA expression in murine PDAC cells harboring wild type p53 (PPT-5123), mutant p53 (PPT-5436) and deleted p53 (PPT-6554 and PPT-W22). Cells were treated for 6h with 12.5μg/ml Etoposide, 3μM AI-10-49 and DMSO as vehicle control. Actin served as housekeeping gene. **E)** Western blot of RUNX1 and p63α upon 6, 24 and 48h treatment with 3μM of AI-10-49 in parental and *NOXA*^*ko*^ MiaPaCa-2 cells and parental PSN1 cells. GAPDH served as loading control. **F)** AnnexinV/PI FACS of parental and *NOXA*^*ko*^ MiaPaCa-2 cells upon 24h treatment with 3μM AI-10-49 and 50μM zVAD-FMK. Upper panel: AnnexinV^+^/PI^−^ fraction. Lower panel: Annexin V^+^/PI^+^ fraction. **G)** Western blot of NOXA protein in MiaPaCa-2 cells. Vinculin served as loading control. **H)** Representative image of clonogenic assay in MiaPaCa-2 parental and NOXA-CRISPR-dCas9-VP64-MS2-HSF1-mediated activation (*NOXA*-CRISPRa). Cells were treated for 3 weeks with vehicle (DMSO) or 400 nM AI-10-49. n=4; all biological replicates were performed as technical triplicates. **I)** Quantification of clonogenic assay in parental and NOXA-CRISPRa. Each treatment was quantified and normalized against its DMSO control. Depicted is the number of colonies in % per treatment compared to vehicle. P value of ANOVA test ***p<0.001.

**SFigure 4** *RUNX1 is upregulated in pancreatic cancer and suppresses apoptosis via NOXA repression*

**A)** Relative *RUNX1* and *NOXA* expression in MiaPaCa-2, Panc-1 and AsPC1 cells upon treatment with a pool of specific siRNAs targeting RUNX1. Cells were treated for 72h with either scrambled siRNAs (Ctrl) or *RUNX1* specific siRNAs. Actin was used as housekeeping gene (ΔΔCt). **B)** GI50 values generated in viability assays using CellTiterGlo in parental, RUNX1ko and cells, determined 72h after AI-10-49 treatment. In addition cells were treated 48h with siRNA (Ctrl and *RUNX1*) and treated for additional 72 h with AI-10-49. **C)** *RUNX1* mRNA expression of patient samples from normal pancreas and pancreatic cancer GEO accession no. GSE15471, GSE16515 **D)** Pearson correlation (incl. linear regression with 95% confidence bands) of PDAC patient samples from the squamous subtype (6) and from microdissected PDAC patient samples (8). **E)** Western blot of RUNX1 and NOXA in parental and *RUNX1*^*ko*^ MiaPaCa-2 cells. *RUNX1*^*ko*^ cells were stably transduced with either a control shRNA or a *NOXA* specific shRNA. Actin served as loading control. **F)** Annexin V FACS analysis of cells as indicated in E).

**SFigure 5** *Quality assessment of H3K27ac ChIP-seq*.

**A)** Fingerprint plot of all replicates including the input controls. Samples with uniform distribution across the genome (e.g. input controls) should plot across the diagonal line, while samples with enrichment across small genomic regions show a steep rise towards the end of the plot. **B)** Correlation plot calculated using the Spearman coefficient quantified by the horizontal bar at the bottom of the plot. **C)** Average signal profile across peaks calculated with macs2 callpeak. The profile was generated with ChIPQC.

**SFigure 6** *RUNX1 suppresses NOXA via the dBS1 downstream enhancer in PDAC cells*

**A)** Chromosome conformation capture (4C, spatial chromatin organisation) and ChIPseq analysis for RUNX1 in indicated cell lines. MiaPaCa-2 cells were treated with vehicle (DMSO)- or 3μM AI-10-49 for 6h. RUNX1 ChIPseq of MCF10A breast epithelial cells (GSE121370) and K562 CML cells (GSE96253) are displayed. The *NOXA* downstream binding site 1 (dBS1) is depicted. **B)** Heatmap of Hi-C data from K562 and MCF10A cells accessed via http://3dgenome.fsm.northwestern.edu/ displaying spatial chromatin organization. **C)** Relative *NOXA* mRNA expression in MiaPaCa-2 and PSN1 cells. Cells were treated with 4μM Entinostat, 10μM Merck60 and 10μM RGFP966 for 24h. Expression was determined by ΔΔCt method. Actin served as housekeeping control. **D)** ChIP-qPCR analysis at the NOXA promoter for H3K27ac and IgG in DMSO- or Merck60 treated MiaPaCa-2 cells. **E)** Western Blot of HDAC1, HDAC2 and HDAC3 in murine PDAC cells harboring a dual recombination system. 600 nM 4-Hydroxytamoxinfen (4-OHT) treatment induced CreERT2 shuttling and excision of either Hdac1 (PPT-F3641) or Hdac2 (PPT-F1648). Actin served as loading control. **F)** Relative *Noxa* mRNA expression in murine PPT-F3641 and PPT-F1648 cells. Cells were treated with 600nM 4-OHT for 48h. Actin served as housekeeping control. **G)** Sequence analysis using ConTra v3 to identify conserved RUNX1 consensus sequences (as indicated) in the dBS1 region. **H)** Upper Panel: Scheme of the human dBS1 region (parental and knockout) with indicated primer binding sites. Lower Panel: Genotyping PCR of MiaPaCa-2 cells to screen for a dBS1 knockout. wt: wildtype; ko: knockout.

**STable 1** *Mean fold change of drug hits from drug screening experiment*

**STable 2** *SwissTargetPrediction of AI-10-49*

**STable 3** *Enriched gene sets of AI-10-49 versus vehicle control treated MiaPaCa-2 cells*

**STable 4** *PDO Gene Expression Matrix*

**STable 5** *Primer sequences, sgRNAs, Antibodies, murine cell lines*

## References

1. R. L. Siegel, K. D. Miller, A. Jemal, Cancer statistics, 2020. CA Cancer J Clin 70, 7–30 (2020).

2. D. Hanahan, R. A. Weinberg, Hallmarks of cancer: the next generation. Cell 144, 646–674 (2011).

3. R. Hamacher, R. M. Schmid, D. Saur, G. Schneider, Apoptotic pathways in pancreatic ductal adenocarcinoma. Mol Cancer 7, 64 (2008).

4. J. E. Bradner, D. Hnisz, R. A. Young, Transcriptional Addiction in Cancer. Cell 168, 629–643 (2017).

5. Y. Cheng et al., Targeting epigenetic regulators for cancer therapy: mechanisms and advances in clinical trials. Signal Transduct Target Ther 4, 62 (2019).

6. P. Bailey et al., Genomic analyses identify molecular subtypes of pancreatic cancer. Nature 531, 47–52 (2016).

7. E. A. Collisson, P. Bailey, D. K. Chang, A. V. Biankin, Molecular subtypes of pancreatic cancer. Nat Rev Gastroenterol Hepatol 16, 207–220 (2019).

8. E. A. Collisson et al., Subtypes of pancreatic ductal adenocarcinoma and their differing responses to therapy. Nat Med 17, 500–503 (2011).

9. R. A. Moffitt et al., Virtual microdissection identifies distinct tumor- and stroma-specific subtypes of pancreatic ductal adenocarcinoma. Nat Genet 47, 1168–1178 (2015).

10. A. Strasser, S. Cory, J. M. Adams, Deciphering the rules of programmed cell death to improve therapy of cancer and other diseases. EMBO J 30, 3667–3683 (2011).

11. P. Fritsche et al., HDAC2 mediates therapeutic resistance of pancreatic cancer cells via the BH3-only protein NOXA. Gut 58, 1399–1409 (2009).

12. M. Wirth et al., MYC and EGR1 synergize to trigger tumor cell death by controlling NOXA and BIM transcription upon treatment with the proteasome inhibitor bortezomib. Nucleic Acids Res 42, 10433–10447 (2014).

13. K. Lankes et al., Targeting the ubiquitin-proteasome system in a pancreatic cancer subtype with hyperactive MYC. Mol Oncol 14, 3048–3064 (2020).

14. E. Oda et al., Noxa, a BH3-only member of the Bcl-2 family and candidate mediator of p53-induced apoptosis. Science 288, 1053–1058 (2000).

15. C. Ploner, R. Kofler, A. Villunger, Noxa: at the tip of the balance between life and death. Oncogene 27 Suppl 1, S84–92 (2008).

16. L. S. Chuang, K. Ito, Y. Ito, RUNX family: Regulation and diversification of roles through interacting proteins. Int J Cancer 132, 1260–1271 (2013).

17. B. A. Otalora-Otalora, B. Henriquez, L. Lopez-Kleine, A. Rojas, RUNX family: Oncogenes or tumor suppressors (Review). Oncol Rep 42, 3–19 (2019).

18. M. Ichikawa et al., AML-1 is required for megakaryocytic maturation and lymphocytic differentiation, but not for maintenance of hematopoietic stem cells in adult hematopoiesis. Nat Med 10, 299–304 (2004).

19. Y. Lou et al., A Runx2 threshold for the cleidocranial dysplasia phenotype. Hum Mol Genet 18, 556–568 (2009).

20. H. Fukamachi, K. Ito, Growth regulation of gastric epithelial cells by Runx3. Oncogene 23, 4330–4335 (2004).

21. K. Inoue et al., Runx3 controls the axonal projection of proprioceptive dorsal root ganglion neurons. Nat Neurosci 5, 946–954 (2002).

22. R. Mevel, J. E. Draper, A. L. M. Lie, V. Kouskoff, G. Lacaud, RUNX transcription factors: orchestrators of development. Development 146 (2019).

23. M. Uhlen et al., A pathology atlas of the human cancer transcriptome. Science 357 (2017).

24. K. Shigesada, B. van de Sluis, P. P. Liu, Mechanism of leukemogenesis by the inv(16) chimeric gene CBFB/PEBP2B-MHY11. Oncogene 23, 4297–4307 (2004).

25. C. J. Scheitz, T. S. Lee, D. J. McDermitt, T. Tumbar, Defining a tissue stem cell-driven Runx1/Stat3 signalling axis in epithelial cancer. EMBO J 31, 4124–4139 (2012).

26. D. J. Birnbaum et al., Expression of Genes with Copy Number Alterations and Survival of Patients with Pancreatic Adenocarcinoma. Cancer Genomics Proteomics 13, 191–200 (2016).

27. C. J. Scheitz, T. Tumbar, New insights into the role of Runx1 in epithelial stem cell biology and pathology. J Cell Biochem 114, 985–993 (2013).

28. I. Kitabayashi, A. Yokoyama, K. Shimizu, M. Ohki, Interaction and functional cooperation of the leukemia-associated factors AML1 and p300 in myeloid cell differentiation. EMBO J 17, 2994–3004 (1998).

29. X. Zhao et al., Methylation of RUNX1 by PRMT1 abrogates SIN3A binding and potentiates its transcriptional activity. Genes Dev 22, 640–653 (2008).

30. Z. Sheng, S. Z. Wang, M. R. Green, Transcription and signalling pathways involved in BCR-ABL-mediated misregulation of 24p3 and 24p3R. EMBO J 28, 866–876 (2009).

31. M. Chan-Seng-Yue et al., Transcription phenotypes of pancreatic cancer are driven by genomic events during tumor evolution. Nat Genet 52, 231–240 (2020).

32. J. A. Pulikkan et al., CBFbeta-SMMHC Inhibition Triggers Apoptosis by Disrupting MYC Chromatin Dynamics in Acute Myeloid Leukemia. Cell 174, 172–186 e121 (2018).

33. A. Illendula et al., Chemical biology. A small-molecule inhibitor of the aberrant transcription factor CBFbeta-SMMHC delays leukemia in mice. Science 347, 779–784 (2015).

34. D. Gfeller et al., SwissTargetPrediction: a web server for target prediction of bioactive small molecules. Nucleic Acids Res 42, W32–38 (2014).

35. D. Wu, T. Ozaki, Y. Yoshihara, N. Kubo, A. Nakagawara, Runt-related transcription factor 1 (RUNX1) stimulates tumor suppressor p53 protein in response to DNA damage through complex formation and acetylation. J Biol Chem 288, 1353–1364 (2013).

36. E. P. Consortium, An integrated encyclopedia of DNA elements in the human genome. Nature 489, 57–74 (2012).

37. M. Orth et al., Pancreatic ductal adenocarcinoma: biological hallmarks, current status, and future perspectives of combined modality treatment approaches. Radiat Oncol 14, 141 (2019).

38. C. Torres, P. J. Grippo, Pancreatic cancer subtypes: a roadmap for precision medicine. Ann Med 50, 277–287 (2018).

39. A. Biederstadt et al., SUMO pathway inhibition targets an aggressive pancreatic cancer subtype. Gut 69, 1472–1482 (2020).

40. R. M. Fritsch, G. Schneider, D. Saur, M. Scheibel, R. M. Schmid, Translational repression of MCL-1 couples stress-induced eIF2 alpha phosphorylation to mitochondrial apoptosis initiation. J Biol Chem 282, 22551–22562 (2007).

41. M. A. Dengler et al., Discrepant NOXA (PMAIP1) transcript and NOXA protein levels: a potential Achilles’ heel in mantle cell lymphoma. Cell Death Dis 5, e1013 (2014).

42. A. Choi et al., RUNX1 is required for oncogenic Myb and Myc enhancer activity in T-cell acute lymphoblastic leukemia. Blood 130, 1722–1733 (2017).

43. Y. Zhou et al., LRG1 promotes proliferation and inhibits apoptosis in colorectal cancer cells via RUNX1 activation. PLoS One 12, e0175122 (2017).

44. S. Liu et al., RUNX1 inhibits proliferation and induces apoptosis of t(8;21) leukemia cells via KLF4-mediated transactivation of P57. Haematologica 104, 1597–1607 (2019).

45. Y. Cheng et al., RUNX1 promote invasiveness in pancreatic ductal adenocarcinoma through regulating miR-93. Oncotarget 8, 99567–99579 (2017).

46. S. H. Wei et al., Inducing apoptosis and enhancing chemosensitivity to gemcitabine via RNA interference targeting Mcl-1 gene in pancreatic carcinoma cell. Cancer Chemother Pharmacol 62, 1055–1064 (2008).

47. A. Kotschy et al., The MCL1 inhibitor S63845 is tolerable and effective in diverse cancer models. Nature 538, 477–482 (2016).

48. L. Castillo et al., MCL-1 antagonism enhances the anti-invasive effects of dasatinib in pancreatic adenocarcinoma. Oncogene 39, 1821–1829 (2020).

49. X. Sun et al., PROTACs: great opportunities for academia and industry. Signal Transduct Target Ther 4, 64 (2019).

50. C. D. Logsdon et al., Molecular profiling of pancreatic adenocarcinoma and chronic pancreatitis identifies multiple genes differentially regulated in pancreatic cancer. Cancer Res 63, 2649–2657 (2003).

51. M. Uhlen et al., Proteomics. Tissue-based map of the human proteome. Science 347, 1260419 (2015).

52. D. Alonso-Curbelo et al., A gene-environment-induced epigenetic program initiates tumorigenesis. Nature 590, 642–648 (2021).

