## Supplementary figures and images for "NOXA expression drives synthetic lethality to RUNX1 inhibition in pancreatic cancer"

### Supplemental Figure 1

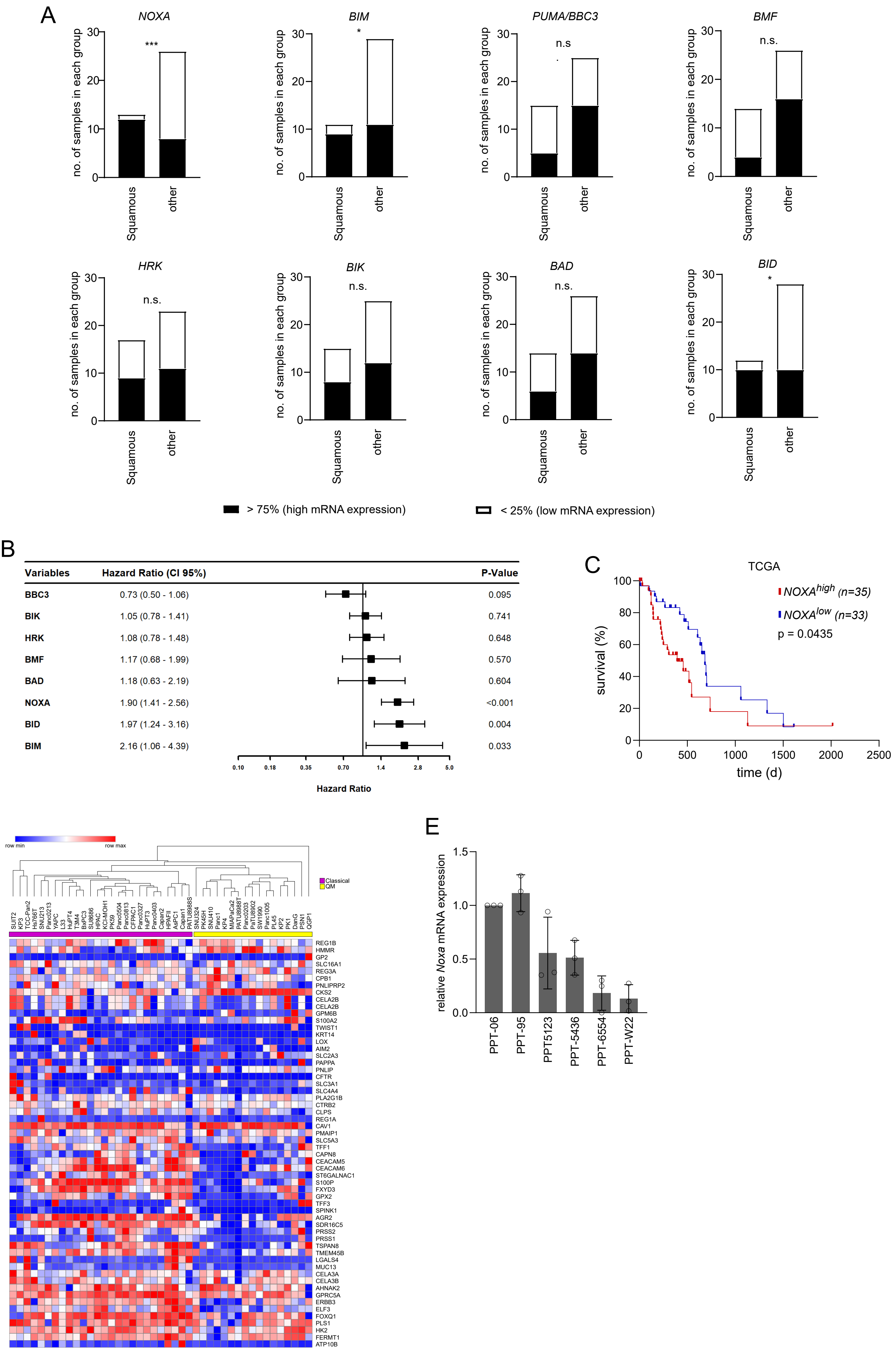

### Supplemental Figure 2

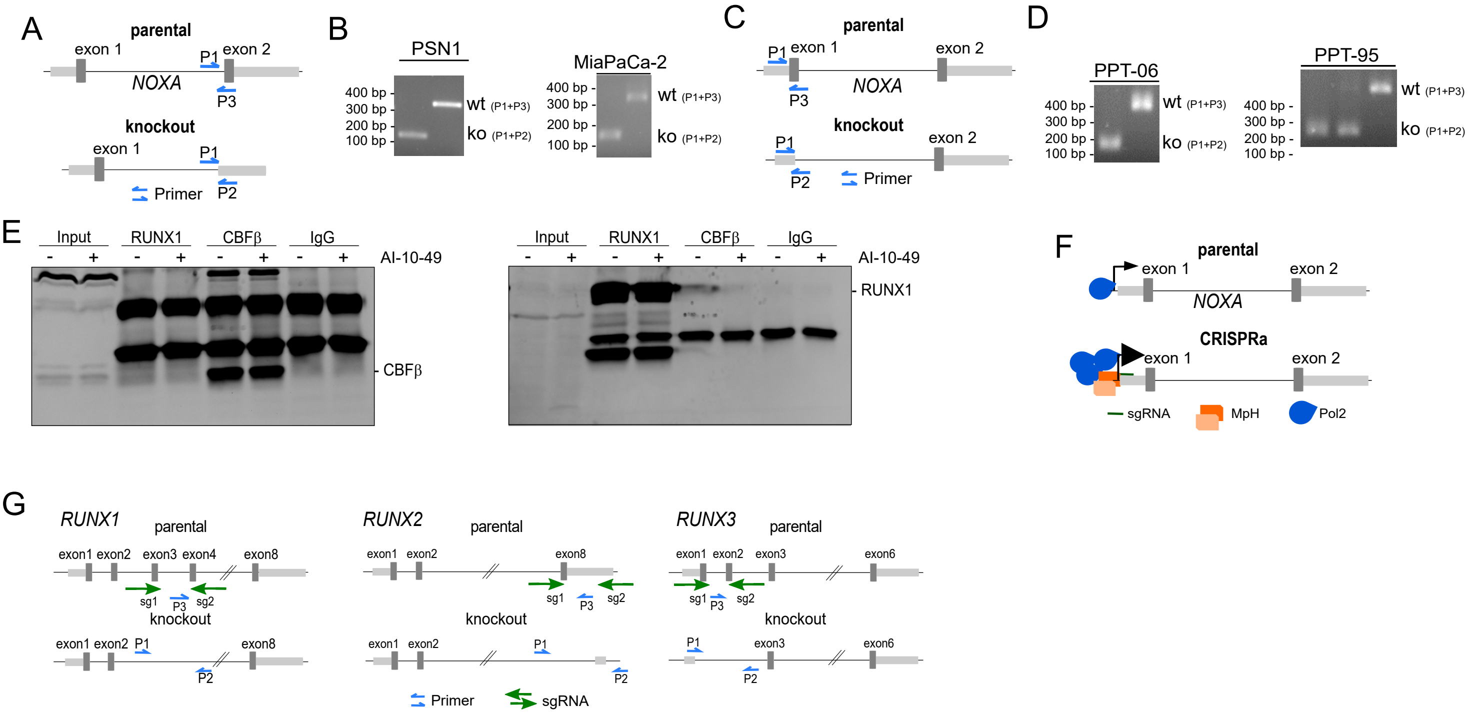

### Supplemental Figure 3

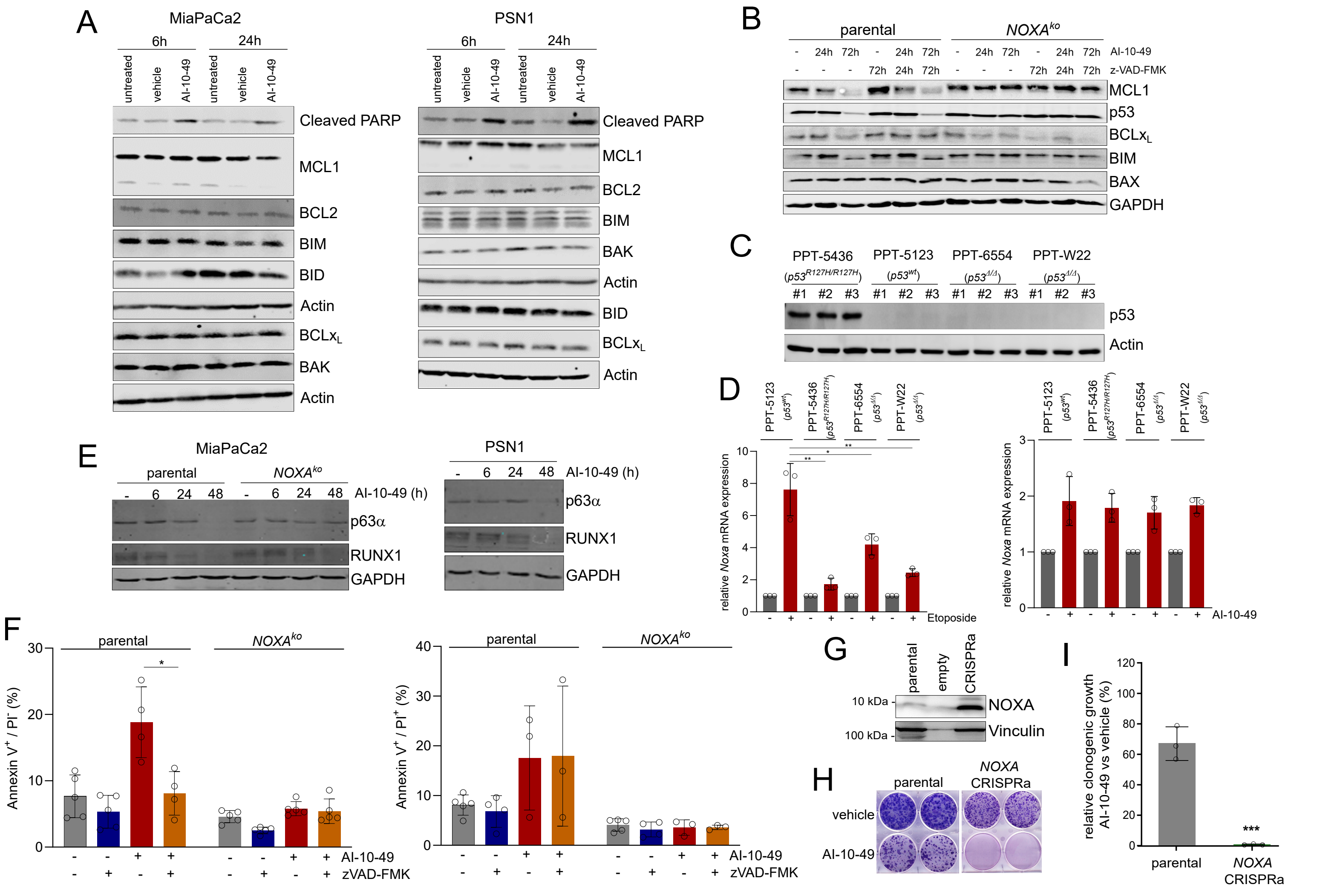

### Supplemental Figure 4

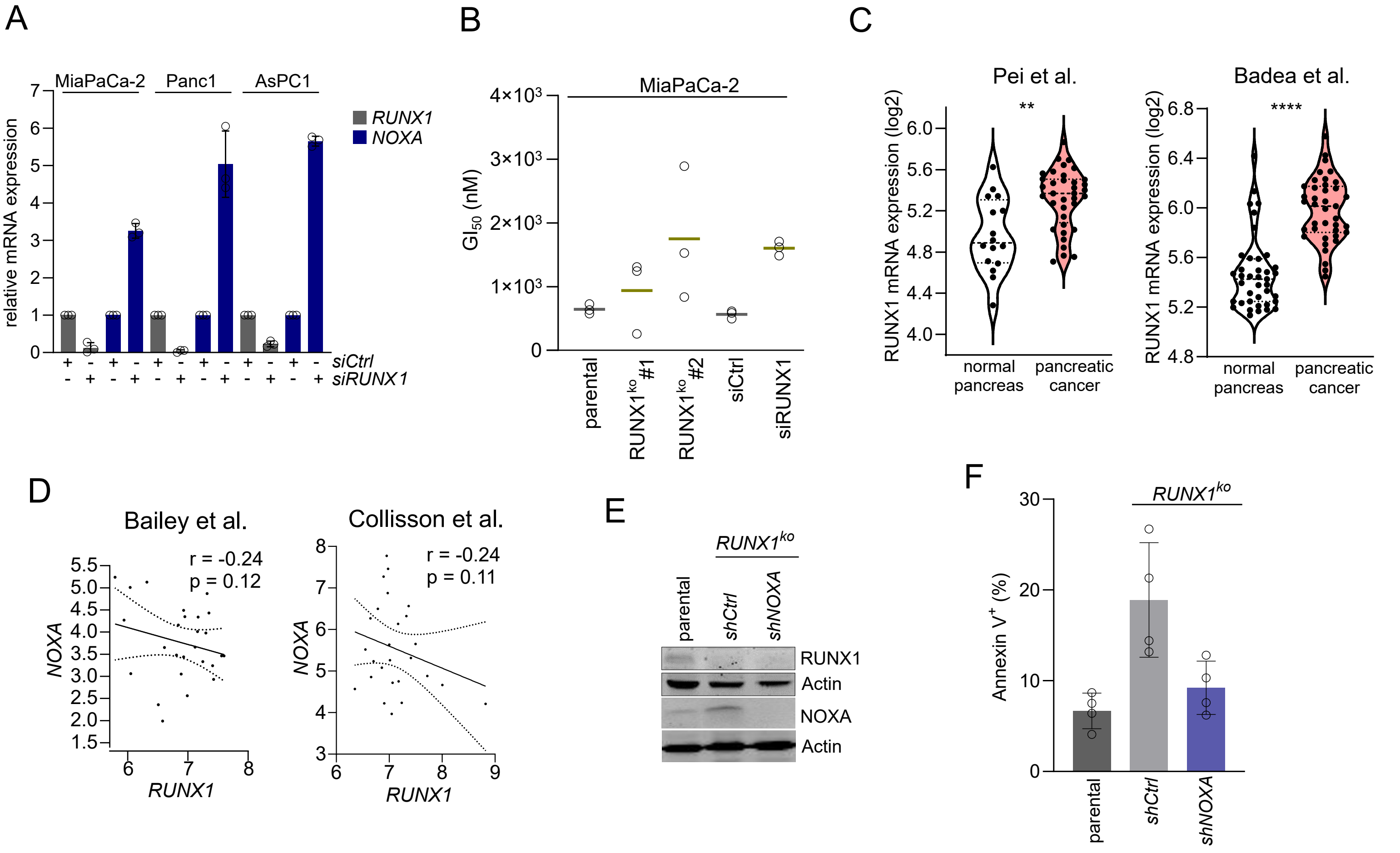

### Supplemental Figure 5

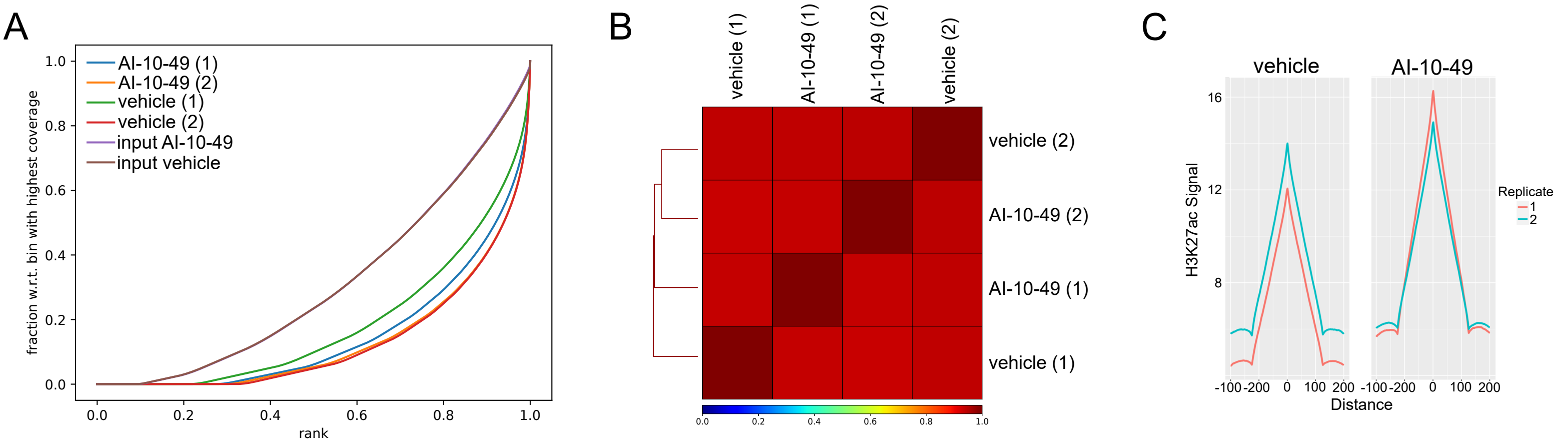

### Supplemental Figure 6

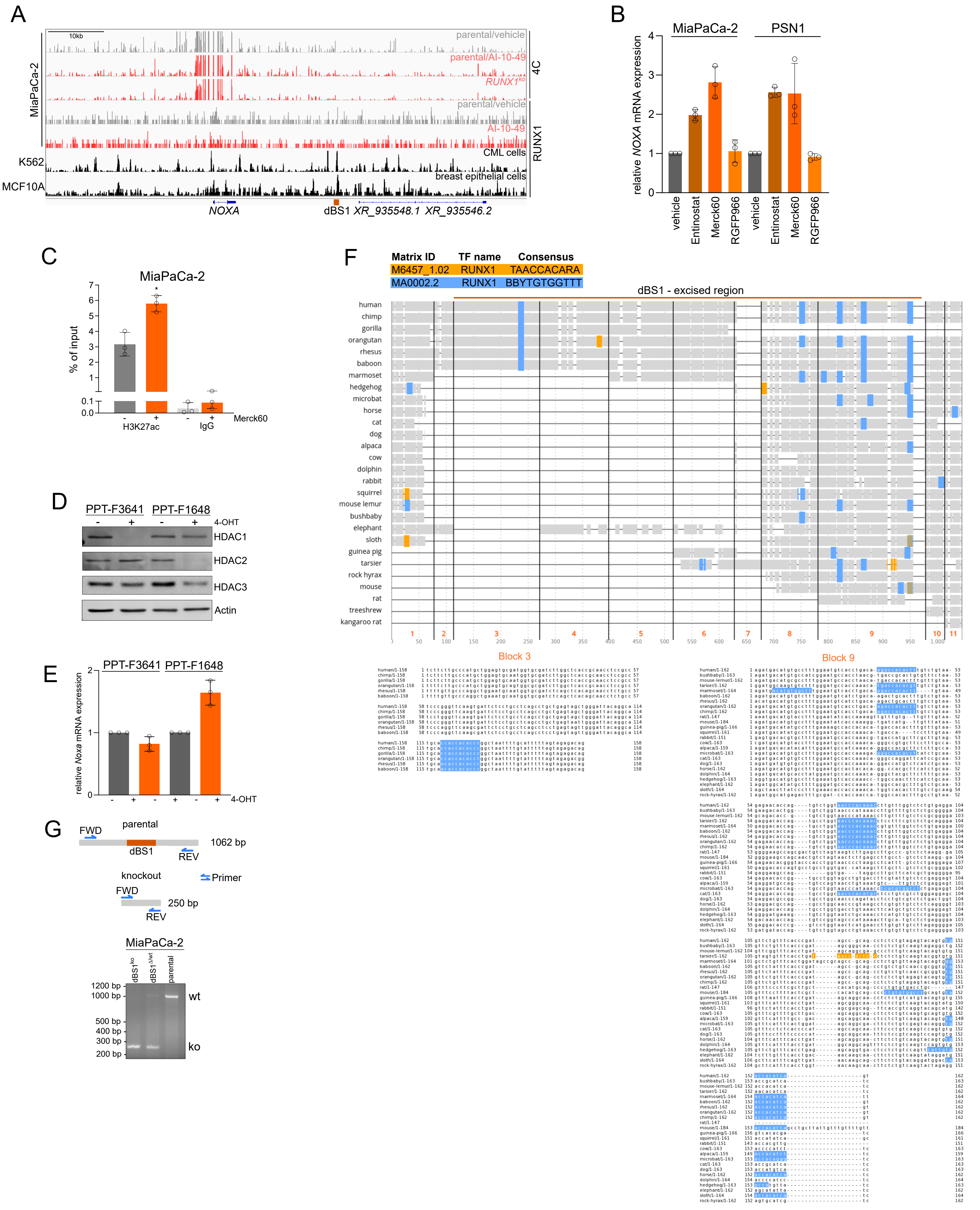
