## Supplemental Material and Methods for "NOXA expression drives synthetic lethality to RUNX1 inhibition in pancreatic cancer"

### Supplementary Materials and Methods

#### Cell culture

Cell lines were cultured in Dulbecco′s Modified Eagle′s Medium (Thermo Fisher Scientific, #41965062) or Roswell Park Memorial Institute (RPMI) 1640 Medium (Thermo Fisher Scientific, #21875091) supplemented with 10% FBS (Thermo Fisher Scientific, #10270106) and 1% P/S (Thermo Fisher Scientific, #15070063). They were passaged up to 14 times in a 1:10 dilution every 3 days. Murine PDAC cell lines were generated from *Kras*$^{G12D}$ driven mouse models as described (1). Genotypes of murine cell lines harboring the dual recominase system (2) - PPT-F1648: *FSF-Kras*$^{G12D/+}$, *FSF-Trp53*$^{del/+}$, *Pdx1-Flp, R26*$^{CAG\text{-}FSF\text{-}CreERT2/+}$, *Pdk1l*$^{oxP/+}$, *Hdac2*$^{loxP/loxP}$; PPT-F3641: *FSF-Kras*$^{G12D/+}$, *FSF-Trp53*$^{del/frt}$, *Pdx1-Flp, R26*$^{CAG\text{-}FSF\text{-}CreERT2/+}$, *Hdac1*$^{loxP/loxP}$, *Hdac2*$^{loxP/+}$, *Hdac3*$^{loxP/+}$. PPT-F1648 and PPT-F3641 cells were treated with indicated concentrations of 4-hydroxytamoxifen (4-OHT) over time to activate the Cre mediated excision of *floxed Hdac* alleles. PCR-based mycoplasma tests (3) were performed at regular intervals.

#### Cell viability assay

Cells were seeded onto 96-well plates (4x10$^3$ cells/well), grown for 24 h and drug-treated. After 72 h of treatment, 10 $\mu$l of MTT reagent (Sigma-Aldrich, #M5655) was added and plates were incubated 4 h at 37°C in the dark. Medium was discarded and MTT crystals were dissolved with 100 µl of a 1:1 DMSO:ethanol solution. Absorbance was measured at 595 nm on a CLARIOstar microplate reader (BMG Labtech, Ortenberg).

#### Drug screening

Two libraries were used for manual high throughput drug screening (FDA approved drug screening library L1300; Cherry pick library L2000, Selleckchem) covering a total of 1842 drugs. The screen was conducted in isogenic human (MiaPaCa-2, PSN1) und murine (PPT-06, PPT-95) cells edited for *NOXA* state. Libraries were provided as individual single-screw-cap compounds in 96-well format dissolved in DMSO at a final concentration of 10 mM. For the drug screening a final concentration of 600 nM was used as previously described(4, 5). Cell viability and readout were done according to previous section. The $OD_{595}$ of vehicle-treated (DMSO) cells was arbitrarily set to 100% and the dose-response was calculated as relative viability. For hit selection, the inhibitors that differentially reduced viability in parental cell lines up to 10% more in comparison to $NOXA^{ko}$ cells were overlapped and those present exclusively in all parental cell lines ($NOXA^{high}$ subtype) were further investigated.

**Dose response assay**

Cells were seeded onto 96-well plates ($4x10^3$ cells/well), grown for 24 h, and then treated with the indicated compounds at seven concentrations (0, 18.75, 37.5, 75, 150, 300, 600 and 1200 nM). Cell viability was analyzed by MTT assay 72 h later. The OD of vehicle-treated (DMSO) cells was arbitrarily set to 100% and the dose-response was calculated as relative viability. GraphPad Prism 8 was used to estimate the half-maximal growth inhibitory concentration ($GI_{50}$) modelled as a non-linear regression and tested for statistical significance by comparing best-fit values of each curve.

**Colony formation assay**

Cells were seeded in 6-well dishes at a density of $2x10^3$ cells/well. After 24 h, medium was exchanged for medium with vehicle (DMSO) or 400 nM AI-10-49 (Biozol, #DL000337516). Medium with drug was refreshed every 3 days. After 2-3 weeks of treatment, the wells were washed with PBS, fixed with methanol for 1 h and stained with Giemsa (Sigma-Aldrich, #GS500) overnight. Plates were scanned and colonies counted with Image J. Percentage of colonies in treatments was normalized against the vehicle control.

**Patient derived organoids culture and viability assay**

PDAC biopsies and tissues were received from endoscopy punctures or surgical resection. 3D organoids were collected, propagated, and analyzed in agreement with the declaration of Helsinki and were approved by the ethical committee of TUM (Project 207/15). Written informed consent from the patients for research use of tumor material was obtained prior to the use. Cellular viability of human patient derived organoids (PDO) was determined using the CellTiter-Glo 3D ATP viability assay according to manufacturer's protocol. Briefly, 1000 cells/well were plated in 80µl of PDO medium [PDO medium components: AdDMEM/F12 medium (Life Technologies) supplemented with 10mM HEPES (Life Technologies), 1x GlutaMax (Life Technologies), 1x B27 (Life Technologies), 100µg/ml Primocin (InvivoGen), 1.25 mM N-acetyl-L-cysteine (MilliporeSigma), 100ng/ml rcWnt3a protein (R&D Systems), 500ng/ml rcR-Spondin 1 protein (R&D Systems), 100 ng/mL mNoggin (Preprotech), 50 ng/mL EGF (Life Technologies), 10 nM Gastrin (MilliporeSigma), 100 ng/mL FGF10 (Preprotech), 10 mM Nicotinamide (MilliporeSigma), 10 µM Y-27632 (MilliporeSigma) and 0.5 µM A83- 01 (Tocris)] and 10µl Matrigel (Corning) in a Matrigel:PBS-coated (ratio 1:3 – 1:4) 96 well plate (Corning, Kat.-Nr. 3610) as described(6). 24h after plating of PDOs, a 7-point dilution of AI-10-49 was added and viability was determined by measuring luminescence FLUOstar microplate reader (BMG Labtech, Ortenberg), after 3 days of treatment.

**Western blotting**

Cells were seeded at $2x10^6$ density in 10 cm dishes. The following day, dishes were washed with PBS and medium replaced with medium plus corresponding treatment. At the indicated time points cells were harvested in RIPA buffer with the final concentration: 150 mM NaCl, 1% IGEPAL CA-630, 0.5% sodium deoxycholate (DOC), 0.1% SDS, 50 mM Tris, pH 8.0. For protein lysis, RIPA buffer was supplemented with 1X protease inhibitor (1 pill of Roche cOmplete™, Mini, EDTA-free Protease Inhibitor Cocktail in 10 ml of buffer) and 1X phosphatase inhibitors (1 pill of Sigma PhosSTOP™ in 10 ml of buffer). Protein concentration was determined by Protein Assay (Bio-Rad, #500-0006) with BSA as a standard. Absorbance was read at 595 nm on a CLARIOstar microplate reader (BMG Labtech, Ortenberg).

Proteins were resolved by SDS-PAGE using gradient gel (10-20%), transferred to PVDF membrane 0.2 µm (Thermo Fisher Scientific) and incubated with specific primary antibodies at 4°C overnight. For ECL measurements, membranes were incubated for 1 h at room temperature with HRP-conjugated secondary antibodies and Kit ECL Prime detection reagent (GE Healthcare, #RPN2232) was used as HRP substrate. For ECL visualization ChemoStar PLUS Imager (Intas, science imaging) was used. Housekeeping antibodies were visualized with ChemoStar PLUS Imager with corresponding fluorescent conjugated antibodies. Quantification was done with Image J software.

**Co-Immunoprecipitation**

Cell lysates were generated as described in the section “Western Blotting” and subsequently precleared with protein G magnetic beads (Cell Signaling) for 60 min at 4°C and then incubated with 4µg RUNX1 (Abcam, STable 5), 4µg CBFb (Invitrogen, STable 5) or 4µg IgG (Cell Signaling, STable 5) for 2h. Subsequently the cell lysate / antibody mixture was incubated with protein G magnetic beads (Cell Signaling) at 4°C for additional 1 h. Beads were washed for four times with RIPA buffer and boiled in 2x laemmli buffer for 5 min before western blotting.

**Apoptosis assay by FACS**

For detection of apoptotic cell death, cells were stained with Annexin V/PI by fllow cytometry. Briefly, $1x10^6$ cells were centrifuged at 300 g for 5 min, resuspended in 300 µl 1X Annexin V binding buffer (BD Pharmingen, #8008652), added 4 µl Alexa Fluor®647Annexin V (Biolegend, #640912) and 10 µg/ml PI (Sigma, P4864) and incubated for 15 min at 4°C in the dark. Cells were analyzed by flow cytometry with Cytoflex S (Beckman Coulter). Annexin V staining was quantified as early apoptotic (Annexin V+, PI-) and apoptotic-dead cells (Annexin V+, PI+). Analysis was performed using FlowJo v10.6.0.

**Growth curves by Cell-Live imaging**

Confluency was determined by monitoring cells in real time with a confluency image mask, which was filtered for each cell line specifically. Cell counts were quantified by 2019B Rev2 version of the Incucyte S3 software.

**sgRNA design and CRISPR/Cas9 mediated knockout**

Single guide RNAs (sgRNAs) were designed by means of the online tool Benchling retrieved from https://benchling.com. The most efficient guide combination was picked according to the scoring system previously described(7). Cloning of sgRNAs was done into a modified px330 plasmid kindly provided by Prof. Rada-Iglesias lab. The sgRNAs were tested in pairs and the most efficient combination was kept for further knockout. 1 µg in total of plasmids was transfected into target cells with Lipofectamine 2000 (Thermo Fisher Scientific) and efficiency was assessed 24 h later by GFP signal. Presence of the knockout was assessed in the bulk via PCR and cells were cultured as single cell in 96-well plate. Single cells were left to grow for approximately 2 weeks until a visible colony was formed and screened via PCR for knockout/wildtype with flanking primers. Knockout efficiency was confirmed via western blot. Absence of wildtype allele was assessed by internal controls within the knockout region. Primer sequences are listed in supplementary table 5.

**Lentivirus production CRISPR mediated overexpression**

Lentivirus production was performed as described by Joung and collaborators (8). Briefly, $3x10^{6}$ HEK293T cells were seeded in 10 cm dishes at day 0. At day 1 cells were transfected as described previously (8). Virus was collected and filtered through 0.45 µm membrane every 12 h after 24 h of transfection for 3 days. Spinfection was performed at 32°C, 1000 g for 2 h with a final concentration of $5x10^{5}$ cells and 8 µg/ml polybrene. CRISPR stable overexpression was performed with lenti dCAS9-VP64_Blast,lenti MS2-P65_HSF1_Hygro and the sgRNA was cloned in lenti sgRNA(MS2)_puro backbone as described previously (9). lenti dCAS-VP64_Blast was a gift from Feng Zhang (Addgene plasmid #61425); lenti sgRNA(MS2)_puro backbone was a gift from Feng Zhang (Addgene plasmid #73795 ; http://n2t.net/addgene:73795 ; RRID:Addgene_73795) and lenti MS2-P65-HSF1_Hygro was a gift from Feng Zhang (Addgene plasmid #61426; http://n2t.net/addgene:61426; RRID:Addgene_61426). sgRNA was designed as described in the previous section. Cells were selected with appropriate antibiotic 24 h after spinfection until no viable cells remained in the negative control. Presence of the transgene was assessed with primers listed in supplementary table 5.

**RNA interference**

For siRNA experiments, endoribonuclease-prepared siRNAs (esiRNAs) against RUNX1 (Sigma Aldrich, EHU124231) and against EGFP (Sigma Aldrich, EHUEGFP) were used as controls. The siRNAs were introduced into MiaPaCa-2, Panc1, and AsPC1 cells with Lipofectamine 2000 according to the manufacturer's recommendation. To generate stably expressing MiaPaCa-2 *RUNX1*$^{ko}$ cells with shRNAs against NOXA and a control shRNA, the lentiviral shRNA constructs were introduced according to the lentivirus-based transduction described above. The shRNA constructs were a gift from Georg Häcker and have been previously published (10).

**RNA isolation and quantitative RT-PCR**

Cells were seeded at day 0 in 6-well dishes at a density of $5x10^5$ cells/well. At day 1 cells were treated with vehicle (DMSO) or AI-10-49 (3 μM) for 6 h. Cells were lysed and RNA isolated with Qiagen RNeasy Isolation kit. 1 μg of RNA was retro transcribed with MMLV kit according to manufacturer's recommendations and 10 ng of random primers. Quantitative PCR was performed with 5 nM primers diluted in 20 μl final volume of SYBR Green Standard Buffer 2X (Thermo Fisher Scientific, #K0223). qRT-PCRs were done as follows: 95°C for 10 min, 40 cycles of 95°C for 15 s and 60°C for 1 min and a final melting curve stage in StepOne Plus System (Applied Biosystems).

Expression levels of target genes were determined with the $2^{-\Delta\Delta CT}$ method from technical triplicates and normalized to *ACTB* mRNA. Relative expression (fold change) was calculated with control samples arbitrarily set as 1.

***In vivo* drug efficacy analysis in mice**

Xenograft assays were performed by EPO (Experimental Pharmacology and Oncology, Berlin-Buch). All animal experiments were approved by the local responsible authorities and performed in accordance with the German Animal Protection law. Subcutaneous MiaPaCa-2 xenograft experiments were performed in *NSG* (*NOD.Cg-Prkdcscid Il2rgtm1Wjl/SzJ, NOD scid gamma*) mice. Tumor growth was monitored daily by measurement with a caliper. Once the tumor volume reached 0.2 $cm^3$ mice were treated with vehicle (2% DMSO+30% PEG300+5% Tween 80+ddH2O) or AI-10-49 (200mg/kg daily) for 9 days. Xenograft tumors were harvested, formalin fixed and paraffin embedded after a total of 12 days. Immunohistochemistry (IHC) of tumor sections was performed on 2 μm-thick paraffin sections. IHC analysis was performed using a Ki67 antibody (DakoCytomation, Glostrup, Denmark; 1:50) and a cleaved Caspase 3 (CC3) antibody (Cell Signaling, 9661, 1:100). Slides were scanned using the Aperio digital whole slide imaging (Leica Biosystems, Germany). Analysis of positive stained cells has been performed using the Positive Pixel Count algorithm (standard settings with col. sat. threshold = 0.15) to quantify the amount of a specific stain present in the respective scanned slide images.

**Quantitative Chromatin immunoprecipitation (qChIP)**

MiaPaCa-2 cells were seeded in 15 cm dishes at a density of $3x10^6$. After 24 h, they were treated with vehicle (DMSO) or AI-10-49 (3 μM) for 6 h. Cells were cross-linked in 1% formaldehyde for 10 min and quenched in 0,125 M Glycine for 20 min. Posterior lysis and harvesting was performed using SimpleChIP Enzymatic Chromatin IP Kit (Cell Signaling Technology, #9003) according to manufacturer’s recommendations. 1-10 μg of antibodies of interest were used to pull down the chromatin (Supplementary Table 4) Immunoprecipitated DNA was analyzed by qPCR on a StepOne Plus as described (11). Chromatin occupancy (ChIP-qPCR) for each sample was calculated as % of input.

**Chromatin immunoprecipitation DNA-Sequencing (ChIPseq) analysis and identification of conserved RUNX1 binding sites**

Cells were prepared as described for qChIP. 10 ng of immunoprecipitated DNA was used for library preparation. ChIP-Sequencing was performed on an Illumina HiSeq2500 sequencer after sample preparation with the ThruPLEX V2 DNA protocol (Takara Bio). The targeting depth was set to 20 million reads. The generated reads were single-end unstranded reads with 50 bp length. FastQC was used for quality inspection of the raw sequencing files (v0.11.9, http://www.bioinformatics.bbsrc.ac.uk/projects/fastqc). The bbduk.sh script (BBTools, https://sourceforge.net/projects/bbmap/) was used for adapter clipping at the 3'-end of the reads with the “adapter.fa” file (included in the BBTools suite) providing adapter sequences. The same tool was used for quality trimming of both the 5'- and 3'-end (k=23, mink=11, hdist=1, qtrim=rl, trimq=10, minlen=30). Alignment with bowtie2 (v2.4.4) (12) to the GRCh38 reference genome (Ensembl release 97, July 2019, primary assembly) followed using default parameters. The resulting output in SAM format was converted to BAM format, sorted and indexed using SAMtools (v1.12) (13). Bigwig coverage files were created using BAMtools from the deepTools2 suite (14) (--normalizeUsing RPKM, --binSize 30, --smoothLength 300, --extendReads 200). The plotFingerprint, as well as the multiBamSummary followed by the plotCorrelation functions from the deepTools suite were run on the sorted BAM files to assess experiment quality (SFig 5A-B)

Prior to peak calling duplicates were filtered out with macs2 filterdup and fragment length was predicted with macs2 predicted to be on average at 301 base pairs in length (15). Peaks were called separately for each replicate using macs2 callpeak with an input control (--nomodel --extsize 301). ENCODE Blacklisted regions (v3) (16, 17) were removed after peak calling from the resulting narrowPeak and treatment pileup bedgraph files. The pileup file was sorted to remove non-primary scaffolds and replicates belonging to the same condition were summarized for visualization purposes with wiggletools mean. The computeMatrix and plotHeatmap tools (deepTools) were then used to create a heatmap around the transcription start sites of all annotated genes.

Sample quality was further assessed with the ChIPQC package using narrowPeak files as input. The DiffBind package was used for differential peak calling (dba.analyze with method=DBA_DESEQ2). Genomic visualizations were performed with pyGenomeTracks tool of the deepTools suite and IGV (18). Note that RUNX1 ChIPseqs were of poor quality, probably due to poor RUNX1 binding or other as yet unidentified factors. Conserved RUNX1 consensus sequences have been analyzed using ConTra v3 (19) using stringency setting of core = 0.9, similarity matrix = 0.75 of the RUNX1 binding motifs of the matrix IDs: MA0002.2 and M6457_1.02.

**Chromosome Conformation Capture Coupled to High-Throughput Sequencing (4C-Seq)**

Cells were were treated with DMSO or AI-10-49 3 μM for 6 h. The crosslinking was performed with 1% of formaldehyde for 20 min and quenched with 0.125 M Glycine for 10 min. The downstream processing of the samples, including the 4C-seq libraries, were generated from cross-linked cells as described previously (20). NlaIII (four-cutter) was used as primary restriction enzyme. DpnII was used as secondary restriction enzyme (four-cutter).

The cross-linked cells were washed with PBS and resuspended in lysis buffer (10 mM NaCl 250 mM Tris HCl (pH 8.0), 0.2% NP-40, 1X Protease inhibitor) for 10 min on ice. Afterwards, cells were centrifuged at 650 g for 5 min at 4°C, the nuclei were re-suspended in 0.5 ml of 1.2X NlaIII buffer and incubated at 37°C and 900 rpm for 1 h. Next, Triton X-100 was added to a final concentration of 2% followed by 1 h incubation at 37°C on a shaker at 900 rpm. The chromatin was then digested with 400 U of NlaIII overnight at 37°C and 900 rpm. NlaIII was inactivated by SDS addition in a final concentration of 1.6% and the mix at 65 °C for 20 min on a shaker at 900 rpm. The digested chromatin was transferred to 50 ml conical tubes and 6.125 ml of 1.15X ligation buffer was added together with Triton X-100 1%. The tubes were incubated on a shaker for 1 h at 37 °C. Subsequently, the digested chromatin was ligated with 100 U of T4 DNA ligase for 8 h at 16 °C followed by RNase A treatment for 45 min at 37 °C. Moreover, the chromatin was treated with 300 mg of Proteinase K and incubated at 65 °C overnight. The resulting DNA was then purified by standard phenol/chloroform extraction, precipitated with ethanol, and re-suspended in 100 ml of ddH2O. At this point the digestion and ligation efficiencies were corroborated by analyzing a small fraction of the purified DNAs by gel electrophoresis as previously indicated (20).The remaining DNA was digested with 50 U of DpnII at 37°C overnight. The resulting DNA was then purified by standard phenol/chloroform extraction, precipitated with ethanol and re-suspended in 500 ml of ddH2O. Afterwards, a second ligation was performed overnight at 16°C by adding 200 U of T4 DNA ligase into a final volume of 14 m 1X ligation buffer (50 mM Tris HCl (pH 7.6), 10 mM MgCl2, 1 mM ATP, 1 mM DTT). DNA samples were again purified by phenol/chloroform extraction, ethanol precipitation and re-suspended in 100 µl dH2O. This product was then purified with the commercial kit QIAgen PCR purification following manufacturer's instructions (Qiagen). For each sample a total of 1 µg of each library was amplified by inverse PCR with the Expand™ Long Template PCR System with 30 amplification cycles (94°C 2 min, 30x [94°C 10s,60°C 1 min, 68°C 3 min], 68°C 5 min). The primers were designed samples specific following the public guidelines (20) targeting the selected viewpoint (NOXA promoter, STable 5). The purified libraries were measured by QuantiFluor® ONE dsDNA System and sequenced on a HiSeq 2500 (Illumina). The processing of the raw sequencing files was identical to the ChIP-Sequencing analysis

**RNA sequencing and analysis**

All samples were sent to Novogene (Cambridge, UK) where they were sequenced on a HiSeq2500 Illumina machine with a target read depth of 20 million reads. The sequencing was paired-end, unstranded, with 150 base pairs per read. The following analyses were performed inside a Bioconda environment (v4.10.1) (21). Quality control was performed using FatsQC. The adapter sequence ("GATCGGAAGAGCACACGTCTGAACTCCAGTCAC"), was removed from the 3'-end of both the forward and the reverse reads using cutadapt (v3.4) (22), followed by quality trimming using Trimmomatic (v0.39, PE mode, MAXINFO:75:0.5) (23). Salmon (v1.5.2) (24) was used for transcript quantification. The reference transcriptome corresponded to GRCh38 (Ensembl release 97, July 2019) (25) with precomputed decoy sequences and was downloaded from a repository maintained by the developers of Salmon (https://drive.google.com/drive/folders/1MeM5BCyUjPD9UTzRUJpAWgI_q4zJj42n) and was additionally augmented with the ncRNA transcriptome from the same Ensembl release. GC-

Bias and sequencing bias were accounted for using the --gcBias and --seqBias flags. The resulting “quant.sf” files were imported into an R (v4.0.4) programming environment and analyzed through an RStudio (v1.4.1106) user interface. A transcript to gene map was created with biomaRt. Tximport was utilized to create a count matrix of isoforms (countsFromAbundance = “dtuScaledTPM”) and genes (countsFromAbundance = “lengthScaledTPM”).

**Differential gene / isoform expression analysis and microarray analysis**

The analyses of differential gene and isoform expression differed only in the input count matrix and were otherwise identical. Normalization was performed with edgeR (v3.32.1) (26). Features (genes or isoforms) with low expression were removed via the filterByExpr function. Afterwards TMM- and library size normalization was performed with calcNormFactors. The model matrix along with the filtered and normalized count table were supplied to the voomWithQualityWeights function of the limma package (v3.46.0) (27-29). Differential expression was then called with lmFit, contrast.fit and eBayes.

Microarray data from 43 PDAC cell lines (Affymetrix, HG-U133-Plus-2.0) from CCLE (30) were processed as shown previously (4, 11). Hierarchical clustering using Pearson's correlation distance/complete linkage and squared Euclidean distance method has been performed using Morpheus (https://software.broadinstitute.org/morpheus/) and ClustVis (31).

**Gene Set Enrichment Analysis (GSEA)**

GSEA was performed using the fgsea (v.1.16.0) package. Genes were ranked according to the t-statistic as a measure of differential expression. The fgsea function was run using default parameters for the hallmark and curated (c2) genesets as defined in the Molecular Signatures Database v7.4 (32, 33). The ClusterProfiler and enrichplot packages were used for visualization purposes.

**Omni-Assay for Transposase-Accesible Chromatin with sequencing (ATAC seq)**

MiaPaCa-2 cells were treated with vehicle (DMSO) or AI-10-49 (3 µM) for 6 h and MiaPaCa-2 *RUNX1*$^{ko}$ cells were harvested in 2 biological replicates. $5x10^4$ cells were collected, washed and snap frozen. Libraries were prepared according to the Omni-ATAC protocol and sequenced on an Illumina HiSeq 2500 sequencer. Paired-end reads were generated of 50 base-pairs in length. The processing of the raw sequencing files was identical to the ChIP-Sequencing analysis up until the creation of a sorted BAM file. The BAM files were then filtered to remove duplicate reads with MarkDuplicates from Picard Tools (v2.26.0). In addition, reads mapping to the mitochondrial genome, with a mapping quality score of <10 or not properly paired were discarded. A bigWig Coverage file was created from the filtered BAM file with the same parameters as in ChIP-Seq.  Peaks were called with macs2 callpeak (--nomodel --shift -100 --extsize 200 --keep-dup all). The mean of replicates belonging to the same condition was calculated after scaling of the pileup file according to library size with wiggletools. A heatmap around transcription start sites was the created as in the ChIP-Seq analysis described above.

**Statistical analysis**

For survival analysis Simple Cox Regression models were constructed using the expression of all "classical BH3 family members". Genes with p-values < 0.1 were used to construct a multiple Cox regression model. For each dataset, normality was assessed by Shapiro-Wilk test and homoscedasticity of the residuals was tested by Levene’s test. Analyses were done with Infostat statistical software or GraphPad Prism v8. For analysis of significant differences, Student’s t-test, analysis of variance (ANOVA) test or Fisher’s exact test was used as indicated in the figure legends. P value < 0.05 was taken as significant. *p<0.05, **p<0.01, ***p<0.001.

#### References (Supplements)


1. J. von Burstin *et al.*, E-cadherin regulates metastasis of pancreatic cancer in vivo and is suppressed by a SNAIL/HDAC1/HDAC2 repressor complex. *Gastroenterology* **137**, 361-371, 371 e361-365 (2009).
2. N. Schonhuber *et al.*, A next-generation dual-recombinase system for time- and host-specific targeting of pancreatic cancer. *Nat Med* **20**, 1340-1347 (2014).
3. J. M. Ossewaarde, A. de Vries, T. Bestebroer, A. F. Angulo, Application of a Mycoplasma group-specific PCR for monitoring decontamination of Mycoplasma-infected Chlamydia sp. strains. *Appl Environ Microbiol* **62**, 328-331 (1996).
4. K. Lankes *et al.*, Targeting the ubiquitin-proteasome system in a pancreatic cancer subtype with hyperactive MYC. *Mol Oncol* **14**, 3048-3064 (2020).
5. C. L. Christensen *et al.*, Targeting transcriptional addictions in small cell lung cancer with a covalent CDK7 inhibitor. *Cancer Cell* **26**, 909-922 (2014).
6. Z. Dantes *et al.*, Implementing cell-free DNA of pancreatic cancer patient-derived organoids for personalized oncology. *JCI Insight* **5** (2020).
7. J. G. Doench *et al.*, Optimized sgRNA design to maximize activity and minimize off-target effects of CRISPR-Cas9. *Nat Biotechnol* **34**, 184-191 (2016).
8. J. Joung *et al.*, Genome-scale CRISPR-Cas9 knockout and transcriptional activation screening. *Nat Protoc* **12**, 828-863 (2017).
9. S. Konermann *et al.*, Genome-scale transcriptional activation by an engineered CRISPR-Cas9 complex. *Nature* **517**, 583-588 (2015).
10. A. Weber, Z. Kirejczyk, S. Potthoff, C. Ploner, G. Hacker, Endogenous Noxa Determines the Strong Proapoptotic Synergism of the BH3-Mimetic ABT-737 with Chemotherapeutic Agents in Human Melanoma Cells. *Transl Oncol* **2**, 73-83 (2009).
11. M. Wirth *et al.*, MYC and EGR1 synergize to trigger tumor cell death by controlling NOXA and BIM transcription upon treatment with the proteasome inhibitor bortezomib. *Nucleic Acids Res* **42**, 10433-10447 (2014).

12. B. Langmead, S. L. Salzberg, Fast gapped-read alignment with Bowtie 2. *Nat Methods* **9**, 357-359 (2012).
13. H. Li *et al.*, The Sequence Alignment/Map format and SAMtools. *Bioinformatics* **25**, 2078-2079 (2009).
14. F. Ramirez *et al.*, deepTools2: a next generation web server for deep-sequencing data analysis. *Nucleic Acids Res* **44**, W160-165 (2016).
15. J. Feng, T. Liu, B. Qin, Y. Zhang, X. S. Liu, Identifying ChIP-seq enrichment using MACS. *Nat Protoc* **7**, 1728-1740 (2012).
16. E. P. Consortium, An integrated encyclopedia of DNA elements in the human genome. *Nature* **489**, 57-74 (2012).
17. H. M. Amemiya, A. Kundaje, A. P. Boyle, The ENCODE Blacklist: Identification of Problematic Regions of the Genome. *Sci Rep* **9**, 9354 (2019).
18. J. T. Robinson *et al.*, Integrative genomics viewer. *Nat Biotechnol* **29**, 24-26 (2011).
19. L. Kreft *et al.*, ConTra v3: a tool to identify transcription factor binding sites across species, update 2017. *Nucleic Acids Res* **45**, W490-W494 (2017).
20. R. W. Brouwer, M. C. van den Hout, I. W. F. van, E. Soler, R. Stadhouders, Unbiased Interrogation of 3D Genome Topology Using Chromosome Conformation Capture Coupled to High-Throughput Sequencing (4C-Seq). *Methods Mol Biol* **1507**, 199-220 (2017).
21. B. Gruning *et al.*, Bioconda: sustainable and comprehensive software distribution for the life sciences. *Nat Methods* **15**, 475-476 (2018).
22. M. MARTIN, Cutadapt removes adapter sequences from high-throughput sequencing reads. *EMBnet.journal* **17**, 10-12 (2011).
23. A. M. Bolger, M. Lohse, B. Usadel, Trimmomatic: a flexible trimmer for Illumina sequence data. *Bioinformatics* **30**, 2114-2120 (2014).
24. R. Patro, G. Duggal, M. I. Love, R. A. Irizarry, C. Kingsford, Salmon provides fast and bias-aware quantification of transcript expression. *Nat Methods* **14**, 417-419 (2017).
25. F. Cunningham *et al.*, Ensembl 2019. *Nucleic Acids Res* **47**, D745-D751 (2019).

26. M. Gierlinski *et al.*, Statistical models for RNA-seq data derived from a two-condition 48-replicate experiment. *Bioinformatics* **31**, 3625-3630 (2015).
27. C. W. Law, Y. Chen, W. Shi, G. K. Smyth, voom: Precision weights unlock linear model analysis tools for RNA-seq read counts. *Genome Biol* **15**, R29 (2014).
28. R. Liu *et al.*, Why weight? Modelling sample and observational level variability improves power in RNA-seq analyses. *Nucleic Acids Res* **43**, e97 (2015).
29. M. E. Ritchie *et al.*, limma powers differential expression analyses for RNA-sequencing and microarray studies. *Nucleic Acids Res* **43**, e47 (2015).
30. J. Barretina *et al.*, The Cancer Cell Line Encyclopedia enables predictive modelling of anticancer drug sensitivity. *Nature* **483**, 603-607 (2012).
31. T. Metsalu, J. Vilo, ClustVis: a web tool for visualizing clustering of multivariate data using Principal Component Analysis and heatmap. *Nucleic Acids Res* **43**, W566-570 (2015).
32. A. Liberzon *et al.*, The Molecular Signatures Database (MSigDB) hallmark gene set collection. *Cell Syst* **1**, 417-425 (2015).
33. A. Subramanian *et al.*, Gene set enrichment analysis: a knowledge-based approach for interpreting genome-wide expression profiles. *Proc Natl Acad Sci U S A* **102**, 15545-15550 (2005).
